# Using Random Forests to Infer Nonlinear Step Selection Effects

**DOI:** 10.1101/2025.03.27.644749

**Authors:** Phillip Koshute, Michael Robinette, William F. Fagan

**Affiliations:** Johns Hopkins Applied Physics Laboratory, 11100 Johns Hopkins Rd., Laurel, MD 20723; University of Maryland, Department of Mathematics, 4176 Campus Dr., College Park, MD 20742; University of Maryland, Department of Biology, 4094 Campus Dr., College Park, MD 20742

**Keywords:** Animal movement, machine learning, partial dependence plots, random forests, step selection

## Abstract

1. Step selection functions (SSFs) examine the factors that motivate animal movement by pairing observed steps with unobserved “comparison steps” and quantifying the effects of covariates corresponding to the values of the factors at each observed step. Traditionally, SSFs are fit with conditional logistic regression (CLR) models, which explicitly reflect how an animal must decide between an observed step and other available steps. These models, however, cannot infer nonlinear effects unless the specific parametric form (e.g., quadratic) of the nonlinear effect is specified.
2. To enable more general inference of nonlinear effects, we propose replacing CLR with random forests (RFs). Using a fitted model, we also describe how to plot the relationship between changes in the value of a covariate upon the likelihood that the animal will select a given step with those covariate values (an “effect curve”).
3. To evaluate our models and their corresponding effect curves, we simulated tracks with various nonlinear effects. RFs yield effect curves that closely resemble the specified nonlinear effect curves, achieving mean R^2^ = 0.953 and outperforming a CLR model with quadratic terms and spline-based generalized additive models. Additionally, we applied our best-performing RF to real wolf data (Barry et al., 2020). The resulting effect curves confirm the authors’ assumptions that their covariates affect wolf movement according to a step function.
4. These results suggest that RFs may be used either to discover unknown nonlinear step selection effects or to confirm previously hypothesized nonlinear effects. Going forward, we envision the use of RFs as an approach that complements existing SSF approaches, providing insights about nonlinear relationships between covariate values and the effects that they have on animal movement decisions.

## 1. Introduction

Ecologists and biologists seek fundamental insights in how animals move through their environment, traveling away from, towards, and with each other (Abrahms et al., 2021; Costa-Pereira et al., 2022]. In particular, conservation biologists aim to anticipate how changes in climate, landscape, or nearby human infrastructure are likely to affect animal movement (Katzner & Arlettaz, 2019). Given an understanding of how animals make movement decisions, landscape accommodations may be implemented to facilitate animal movements.

Others track animal movements with particular applications in mind. Military planners may be informed by the movements of animals both for training exercises (Krausman et al., 2007) and tactics (Smith, 2018). Public health officials monitor the migration, home range shifts, or other movement of animals bearing zoonotic diseases (Reed et al., 2003; Ratanakorn et al., 2012). For them, anticipating the likelihood of overlap between animals’ movements and human activity can enable appropriate preparations (Roden-Reynolds et al., 2022). For these researchers and practitioners, having an understanding of animals’ movement decision-making can enable assessments of those animals’ likely movements, either under current conditions or if the landscape would be altered.

In recent decades, technology for tracking animals and the factors that may affect their movement has increased in capability and accessibility. For instance, animal tracking studies that began with radio collars in the 1960s (Craighead & Craighead, 1972) have capitalized on the increasing availability of GPS-based location tracking and remote reporting (Kays et al., 2015). Devices are fastened on animals and report the animals’ locations at specified time intervals. As such tracking devices decrease in size and weight, the range of animals that can be remotely studied broadens. Likewise, longer battery lives increase the feasibility of longer studies, ultimately enabling the collection of more tracking data and more sophisticated analyses. Wide availability of satellite-measured environmental factors of interest permits animal tracks to be paired with digitally mapped covariates (Kays et al., 2022).

Established terminology describes these animal location data. Pairs of consecutive reported animal locations comprise steps; sequences of steps comprise movement tracks.

These tracks indicate where an animal has gone. On their own, though, tracks do not reveal animals’ motivation for their movement decisions and thus offer limited insights on where an animal is likely to move amid changing conditions.

Step selection functions (SSFs) provide a quantitative framework for analyzing the effect of environmental factors and other step characteristics upon animal movement (Thurfjell et al., 2014). Originally proposed by Fortin et al. (2005), SSFs contrast observed steps (OSs) with nearby and feasible but unobserved “comparison” steps (CSs). The set of an OS and its corresponding CSs is known as a stratum. Some researchers refer to CSs as available steps.

The CSs are sampled from distributions of feasible steps, often expressed in terms of step length (SL) and relative turning angle (RTA). For a given OS, each CS shares the same starting point. By retaining a common step starting point for all steps within a stratum, researchers directly compare the step that an animal selected with other steps that were available, thereby focusing on the factors associated with that selected step. These factors are quantified with covariates such as resource levels, environmental conditions, or distances to relevant landmarks. Figure 1 illustrates how CSs are selected for a given OS.

**Figure 1:**
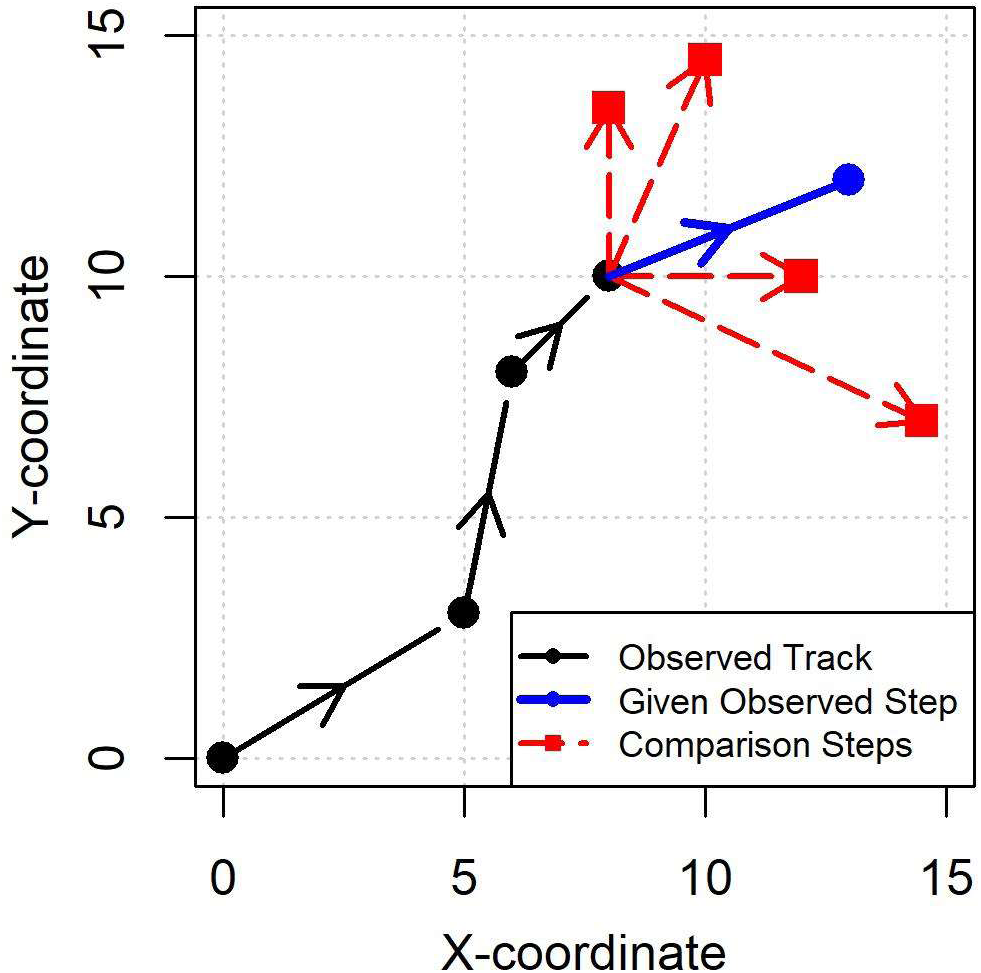
Illustration of Comparison Steps for a Given Observed Step.

The SSF framework enables inference of effects of covariates, traditionally estimated with conditional logistic regression (CLR) (Hosmer et al., 2013). However, effects that can be inferred by CLR are limited to linear effects or specified parametric forms. Thus, this approach has limited ability to infer effects of covariates that are strongest at an intermediate range (Aikens et al., 2017), follow a step function (Barry et al., 2020), or have other nonlinear forms.

Klappstein et al. (2024) recently demonstrated the feasibility of generalized additive models (GAMs) consisting of multiple-degree splines for inferring nonlinear effects, but these methods likewise are limited to specific parametric forms.

Alternatively, models based on machine learning (ML) could infer nonlinear effects without requiring parametric forms to be specified. As one example application of ML methods, we envision adapting random forests (RFs) to infer step selection effects. RFs are collections of decision trees constructed with strategically selected subsets of available data. The results of a given RF are determined by aggregating the results of the decision trees (Breiman, 2001). By aggregating the results of the carefully constructed trees, a RF generally achieves superior performance to the individual trees. Unlike CLR models and GAMs, RFs can infer nonlinear effects without any parametric requirements.

RFs have been previously used as resource selection functions (RSFs) to infer the effects of covariates on mountain lions’ habitat preferences (O’Malley et al., 2024). Unlike SSFs, RSFs do not condition on the starting point of a given step and do not separate steps by strata. Thus, it is natural to apply RFs to RSFs, considering all observed and unobserved locations within a track simultaneously. However, RFs may also be easily applied to SSFs, comparing OSs with CSs conditionally sampled nearby those OSs. As implemented by O’Malley et al. (2024) for RSFs, RFs can yield curves that characterize the effect of each modeled covariate.

Here, we demonstrate the feasibility of RFs for inferring nonlinear effects within animal movement tracks. We follow a simulation-based approach for comparing RFs, CLR models, and GAMs. Simulation offers the key advantage of known specified nonlinear effects against which inferred effects can be compared. We also provide guidance on how to elicit “effect curves” (linear or nonlinear) from RFs. Additionally, we highlight the applicability of RFs to SSFs with data collected from wolves in Finland (Barry, 2020).

## 2. Materials and Methods

Within our study, we follow a rigorous process for comparing model types’ capabilities to infer nonlinear and linear effects. We simulate generic covariate landscapes and tracks within those landscapes. For each OS in each track, we sample feasible CSs. Using the resulting strata of steps, we fit RFs and various baseline models. With a customized approach, we construct a partial dependence plot (PDP) (Friedman, 2000) that illustrates the effect of one or more covariates’ values upon each model for each track. To emphasize how the PDP showcases the varying effects of a covariate, we refer to PDPs in this paper as “effect curves.” We compare the resulting curves with the “true” curve of the specified effect with two different metrics. Additionally, we fit a RF with the best-performing settings to wolf data (Barry, 2020) and show how RFs can inform real-world analyses. We have conducted all analyses using R version 4.4.1 (R Core Team, 2024). All references to packages likewise refer to R. Fig. S-1 depicts how these steps fit together.

### 2.1 Data Simulation

To simulate tracks of animals motivated by specific effects, we simulate landscapes of covariates or resources. See the Supplemental Material for details. Following this process, we simulated five different landscapes, each with three covariates, designated as *F_1_*, *F_2_*, and *F_3_*.

Given a simulated landscape, we simulate a track according to methods used by Avgar et al. (2016) and others. We successively carry out weighted sampling of candidate steps near the track’s current location; for this study, we consider 10000 candidate steps for each step added to the simulated track. We compute the weights of each candidate step from the covariate values at the step.

We randomly sample each candidate step’s SL from an exponential distribution with mean 4 and its RTA from a von Mises distribution with mean 0 and concentration parameter 3. We do not include correlation between SL and RTA.

We do not specify a unit for SL. Instead, SL can be interpreted according to the spatial resolution of the covariate landscape. For instance, SL = 1 corresponds to the width of a single grid within the covariate landscape. We assume that all covariates have the same spatial resolution.

For the three covariates, we specify effects ***β*** that determine the extent to which each covariate affects that weight of each candidate step. We specify different effects for each of the three covariates: no effect (*β_1_* = 0), rendering the first covariate as “noise”; a linear effect (*β_2_* = 2); and an effect for which we specify different linear or nonlinear effects.

We consider four variations of this third effect *β*_3_ (i.e., four “effect types”): linear, ramp-up, step-down, and triangular. These effects dictate the contribution of a given covariate to a selection likelihood or “score” of given candidate step. With a linear effect, the effect is directly proportional to the covariate’s value, e.g., as with certain resources. Traditionally, SSFs (as with CLR) assume linear effects. We opted to include a linear effect in our study to confirm that model types can infer linear effects in addition to nonlinear effects. A ramp-up effect starts at zero but then linearly increases to a certain “saturated” value. For a predator, density of prey may follow a ramp-up effect. A step-down effect is constant up to a certain value before dropping to zero (no effect), as with a resource that can only be perceived within a certain distance. Similarly, Barry et al. (2020) modeled proximity to human infrastructure features according a step-up effect that starts at zero and instantaneously increases to a fixed value beyond a threshold. (The key difference between ramp-up and step-down effects is not their “up” and “down” directions but the steepness of the effect values’ transitions. A ramp-up effect has a gradual transition, while a step-down effect has a sharp transition. We could alternatively have studied this difference with ramp-down and step-up effects.) A triangular effect starts at zero, ramps up to a maximum value, and then ramps back down to zero effect. Having such an intermediate peak may occur when a foraging animal balances quality (low normalized difference vegetation index (NDVI)) with quantity (high NDVI) (Aikens et al., 2017). Figure 2 depicts these four variations and shows the “milestone points” that characterize the shape of each effect and are used to assess fitted effect curves (see Section 2.5). The equations underlying each effect type are given in Table S-1.

**Figure 2:**
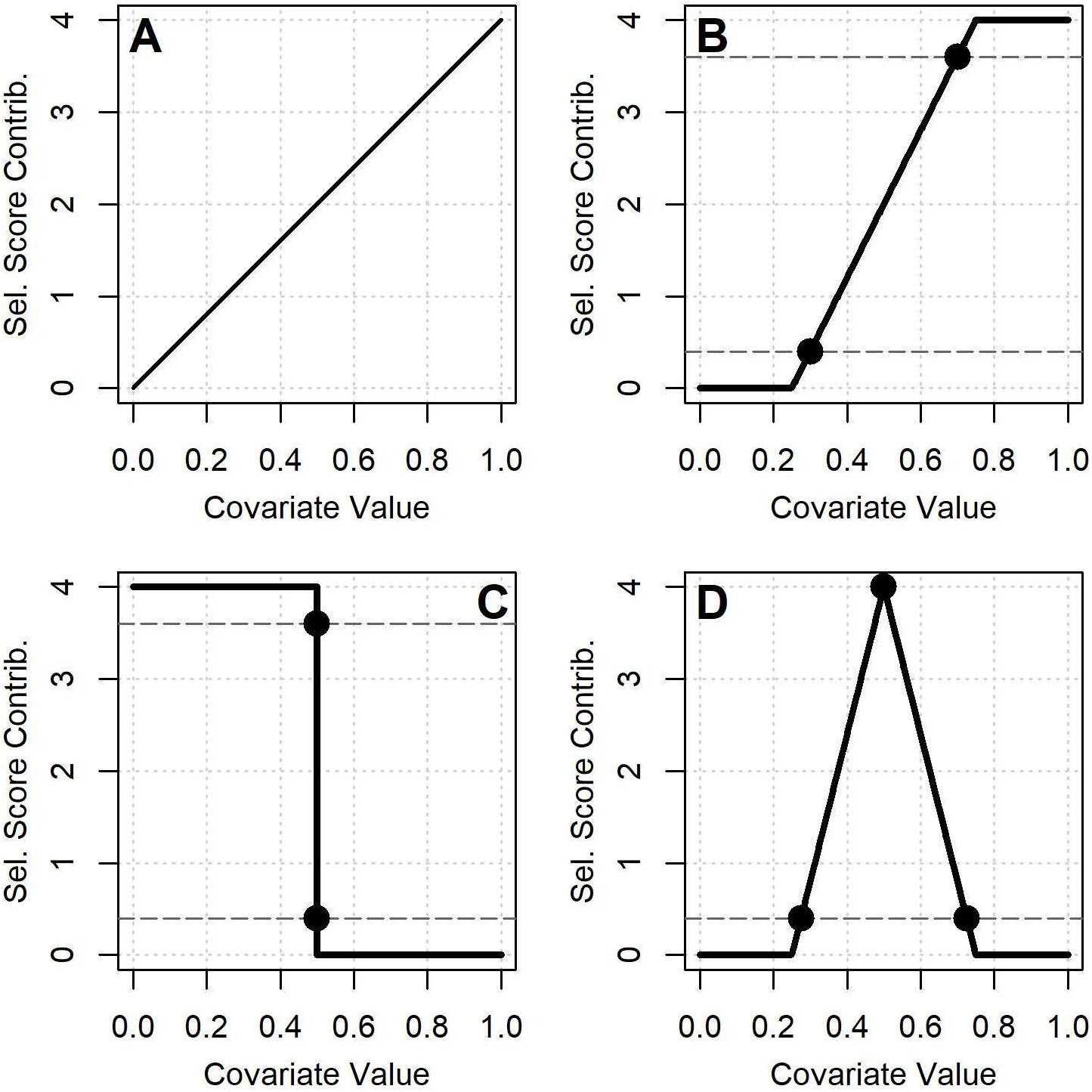
Effect Curves for Different Effect Types: (A) Linear, (B) Ramp-Up, (C) Step-Down, (D) Triangular.

We think of each effect type being associated with an individual animal (each possibly from a different species). Thus, different tracks with the same effect type correspond to different observations of that animal, either in the same landscape or a different landscape.

In general, for the *m*th candidate step 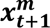 within strata *t*+1, the contributions of these covariates are summarized by the resource-based selection score:

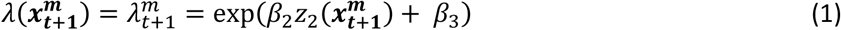

Here, 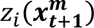 is the value of the *i*th covariate for 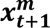 and *β*_3_ is a function of 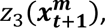 as described in Table S-1. Ultimately, we determine the weights of each candidate step are by this selection probability 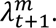

From a given step, we divide the scores of all individual candidate steps by their sum 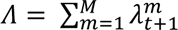 to obtain the weights of each step.

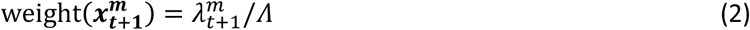

After this scaling, we compute the cumulative weight of the first *m* steps. We uniformly sample a random value between 0 and 1. The candidate step corresponding to the largest cumulative sum less than the random value is the step that is added to the simulated track. In this way, candidate steps with larger weights are more likely but not guaranteed to be sampled.

We continue to iteratively simulate steps according to this process until we have simulated a track with the specified track length *N*. The resulting track has *N*-1 OSs. Because a prior step (involving two prior locations) is required to compute RTA, we use *N*-2 OSs for each model. To coincide with our analyses that involve powers of two, we use *N* = 513, which yields 512 strata per track.

For each of five landscapes and four effect types, we simulated five tracks. Thus, we simulated 100 total tracks.

### 2.2 Comparison Steps

In order to fit the conditional logistic regression models, we sample 32 CSs for each OS. Our approach for sampling CSs is similar to how we sample candidate steps within simulated tracks, except that we do not include covariate-based weighted sampling and we draw SLs and RTAs from fitted rather than specified distributions.

We determine the CSs according to randomly sampled SLs and RTAs. We draw SLs from an exponential distribution with mean *μ_L_*. We draw RTAs from a von Mises distribution with concentration parameter *κ_A_*. We fit both *μ_L_* and *κ_A_* to the observed track, fitting *μ_L_* with the *fitdistr* function from the *MASS* package (Venables & Ripley, 2002) and fitting *κ_A_* with the *lm.circular* function from the *circular* package (Agostinelli & Lund, 2023). Using fitted parameter values (which may differ from the specified values) rather than specified values more closely mirrors how SSFs are used in practice. We do not model correlation between the SLs and RTAs of the CSs.

This sampling approach yields 33 steps per stratum (1 OS and 32 CSs) that are available for fitting a conditional logistic regression model. For some analyses, we only consider a subset of the available CSs. For CLR models and GAMs, CSs enable an approximation of the sum of the probabilities of the full range of feasible steps (“normalization integral”). (See Michelot et al. (2024) for more details on this concept.) For RFs, CSs serve a different purpose, instead providing negative examples to allow the models (as classification models) to find patterns within covariates that distinguish between observed and unobserved steps.

### 2.3 Types of Models

For each number of CSs, we compare 16 types of models: eight RFs, six GAMs, and CLR models with and without quadratic terms. For RFs, we vary the number of steps used to train each tree. For GAMs, we vary the number of degrees of freedom (dfs) for each spline. Following the integrated step selection approach outlined by Avgar et al. (2016), we include SL and the cosine of the RTA as covariates for each model. This amounts to five covariates for each model (*F_1_*, *F_2_*, *F_3,_* SL, and cos(RTA)).

To construct RFs, we use the *randomForest* package in R (Liaw & Wiener, 2002). For those unfamiliar with RFs, we provide an overview in the Supplemental Material. We consider 4, 8, 16, 32, 64, 128, 256, and 512 OSs and the same number of CSs to train each tree. We opted for such stratified sampling, as well as for sampling with replacement based on sensitivity analysis (see Table S-2). For the remaining RF parameters, we use the default *randomForest* options (including 500 trees per forest).

The GAMs and CLR models provide a baseline against which to compare the RFs. For GAMs, we fit models with the *gam* function in the *mgcv* package in R (Wood, 2017). In general, we followed the code provided by Klappstein et al. (2024), varying the number of dfs *k* as 5, 10, 15, 20, 30, or 40. We set *F_1_*, *F_2_*, *F_3_,* as spline-based smooth terms, each with *k* dfs. We specify “cox.ph” as the family to mirror the CLR framework.

Because we are focusing on the effect of individual covariates and we did not model interactions in the simulated tracks, we do not include interaction terms with CLRQ models. Thus, the terms of the CLRQ models are 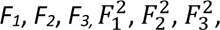 step length, and cos(RTA). Following previous SSF research (e.g., Avgar et al., 2016), we used the *clogit* function within the *survival* package in R (Therneau, 2024) to fit the CLR and CLRQ models. This use leverages the equivalence of the log-likelihood functions for CLR and Cox proportional hazard models (McDonald et al., 2006).

Our CLR models and the GAMs are fit in a way that reflects that exactly one step is selected from each stratum. In contrast, RFs do not inherently reflect the selection of a single step. Nevertheless, CLR models, GAMs, and RFs all ultimately can be interpreted as binary classification models that yield probabilities of the positive class (corresponding to an OS) for a given set of values for the models’ covariates. As described below, we can leverage the variation of this probabilistic output to infer the effect of a covariate of interest.

### 2.4 Inferring Effects

A model’s effect curve describes how a change in covariate values affects the output of the model. While our approach could be extended to multiple-covariate combinations with corresponding higher-dimensional effect curves, we focus on one-covariate effect curves for this demonstration. As examples, Figure 2 above shows idealized versions of effect curves; effect curves fitted to data are discussed in Section 3.

For RFs, such curves can be computed from the *randomForest*::*partialPlot*. However, in order to directly compare effect curves from different types of models, we opted to write custom code. In particular, for a given model’s effect curve, we combine a given *F_3_* value (*z*_3,v_, i.e., the *v*th value of *z*_3_) with many different sets of values for the other covariates (*F_1_*, *F_2_*, SL, and cos(RTA)) and run the resulting combinations through the model, obtaining a predicted probability for each combination. We compute the model’s mean probability *μ*_3,v_ for this *z*_3,v_. For RFs, *μ*_3,v_ corresponds to the proportion of trees predicting the “observed step” class with *F*_3_ = *z*_3,v_. For a given track, we determine the sets of all other variables from all OSs and CSs. For example, with 512 strata per track and 33 steps per stratum, we have 16896 combinations for each value of *F_3_*. We discuss other options for constructing effect curves in the Supplemental Material.

We repeat these computations for 40 different *z*_3,v_, evenly spaced between the 5^th^ and 95^th^ quantile of values of *F_3_* observed within the strata of a given track (including observed and CSs). These *z*_3,v_ do not need to coincide with observed values of *F_3_*. We exclude the lowest and highest 5% of values in order to avoid sparsely represented intervals of values. (This truncation of extreme values would not be possible with *partialPlot*.) The 40 *z*_3_values accompanied by 40 *μ*_3_values from a given model form the basis of that model’s effect curve for *F_3_*.

Rather than directly plotting *z*_3_ versus *μ*_3_, we transform the *μ*_3_ values with half of the logit function, with 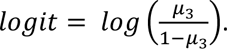 The rationale for this transformation, used also by the *partialPlot* function for plotting effect curves, is outlined by Hastie et al. (2009). For binary classification, transforming a model’s output probability with the half-logit function in order to predict a class minimizes the well-known exponential loss function. Additionally, with two possible classes (as we have with observed and unobserved steps), inserting the transformed value into Equations (1) and (2), e.g., in place of 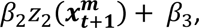 sets each step’s weight to *μ*_3_.

To avoid infinite values from this transformation (i.e., with *μ*_3_= 0 or *μ*_3_= 1), we select a small smoothing constant *C* and convert all values less than *C* to *C* and all values greater than 1-*C* to 1-*C*. All values between *C* and 1-*C* are not affected. For GAMs, we set *C* to *.Machine$double.eps* = 2.2*10^-16^ constant, which is also used by the *partialPlot* function and was confirmed by sensitivity analysis (see Supplemental Material). For RFs, we set *C =* 0.05, also based on sensitivity analysis. Setting a larger value of *C* stretches the intermediate range of the effect curve and downplays the influence of *μ*_3_values close to 0 or 1.

For RFs, the probabilities *μ*_3_depend on the distribution between OSs and CSs within the data. Models fitted to less balanced data (i.e., with more CSs) tend to yield lower *μ*_3_. To account for this phenomenon and to enable comparison across different models, we scale all transformed *μ*_3_values for a given model and track so that they lie within [0, 1]; the lowest *μ*_3_is mapped to 0 and the largest *μ*_3_ is mapped to 1. All intermediate values are scaled linearly between 0 and 1. This scaling is not necessary when constructing an effect curve with a single model.

For easy reference, we provide a summary of the process for constructing an effect curve in the Supplemental Material.

### 2.5 Evaluation Approach

We use two metrics to assess which type of model’s effect curve most accurately capture the specified effect curve: one that considers how well the effect curve inferred from the model matches the entire specified curve (R^2^); and another that considers how well the inferred curve matches the key points or “milestones” of the specified curve (milestone proximity score (MPS)).

R^2^ directly compares the model curve to the specified curve across all values. For a set of values *w_i_* that span the range of the covariate of interest, we compute R^2^ by comparing *w_i_* with the corresponding values *y_i_* in the true effect curve values (which we have specified).

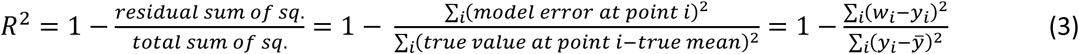

This metric is commonly used to assess how well a model fits a set of data or to compare a model’s predictions with a set of true, labeled data. Higher values of R^2^ indicate inferred effect curves that more closely match the specified effect curve.

As a complement, the MPS is a novel metric that identifies the locations of milestones within the inferred effect curve and assesses their proximity to the locations of milestones in the true effect curve. Milestones include the start and end of the ramp in a ramp-up; the start and end of the decrease in a step-down effect; and the start, peak, and end of the triangular shape in a triangular effect. Milestones for each type of nonlinear effect are shown as circles in Figure 2. More detailed criteria for identifying milestone locations within fitted effect curves are provided in Table S-1. Unlike R^2^, MPS rewards inferred effect curves that have the intended shape but are otherwise shifted or noisy.

To compute the MPS for a fitted and scaled effect curve 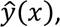 we identify the milestone values for that curve, i.e., the covariate values x when 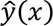 is within a margin *A* of each milestone point. We compute the difference between the fitted curve’s milestone values and the milestone values of the specified effect curve. The overall milestone proximity score for a given effect curve is the mean absolute difference. To evenly compare MPSs between curves, we scale the mean difference by the range of covariate values used to construct the effect curve. Thus, MPS values range from 0 to 1; lower values indicate closer resemblance to the specified effect curve. For instance, MPS = 0.05 indicates that the fitted effect curve’s milestones are on average within 5% of the specified curve’s milestones. For this analysis, we set *A* = 0.1.

We use data from one set of landscapes and tracks (“tuning set”) to determine the best settings for RFs (including number of steps to train each tree and number of CSs), for GAMs (number of dfs and number of CSs), and for the CLR models (number of CSs) in terms of the mean difference of R^2^ minus MPS (not weighting either metric since they vary on roughly the same scale). Given these best settings, we evaluate models with another set of tracks (“test set”), simulated on another set of landscapes, comparing mean R^2^ and MPS. Such an arrangement ensures that we fairly compare RFs with other types of models, removing the potential bias of models fitted to particular tracks.

### 2.6 Real Data

For further demonstration, we fit a RF with the best settings to wolf data (*Canis lupus*) provided by Barry et al. (2020). These data include tracks from 36 wolves in Finland, ranging from 261 to 3919 steps (median = 977 steps), with distance to nearest house, distance to nearest primary road, and distance to nearest forest road as the covariates for each OS and CS (Barry 2020). Prior to fitting SSFs, Barry et al. (2020) converted the continuous values of these covariates to binary covariates, which is equivalent to assuming that the covariates’ effect curves are step functions (e.g., the step-down effect that we considered via simulation). These authors set separate threshold values for each covariate, determining the values by maximizing the model’s likelihood as one covariate was varied at a time. This analysis resulted in thresholds of 600, 600, and 200 m respectively for the house, primary-road, and forest-road covariates.

To examine the authors’ step-function assumption and corresponding thresholds, we fit our RFs to the raw distance values (rather than the binary covariates) and qualitatively compared the resulting effect curves to their findings.

When determining optimal threshold values, Barry et al. (2020) considered all wolves within a single model. Similarly, we fit a single RF to the tracks of all wolves.

## 3. Results

Figure 3 shows the mean R^2^ for different models fitted to the 100 tracks in the tuning set. In this figure, GAMs are labeled as GAM-*VW* and RFs as RF-*XYZ*, with *VW* indicating each spline’s degrees of freedom and *XYZ* indicating the number of OSs used to train each tree. For instance, each tree in a RF-004 model was trained with 4 randomly sampled OSs and 4 randomly sampled CSs (regardless of the number of available CSs). To highlight differences between better values, only R^2^ values between 0.75 and 1 are depicted; values below 0.75 are shown in gray. Darker colors indicate better performance. Table S-5 shows the numerical R^2^ values underlying Figure 3.

The best model is an RF with 512 OSs and 512 CSs used to train each tree (RF-512) with CSs drawn from 8 available CSs per OS (mean R^2^ = 0.951, mean MPS = 0.036). With 4 or more CSs, RF-032, RF-064, RF-128, and RF-512 also have mean R^2^ values that exceed 0.927. Similarly, with any number of CSs, RF-512 has mean R^2^ values that exceed 0.941.

**Figure 3:**
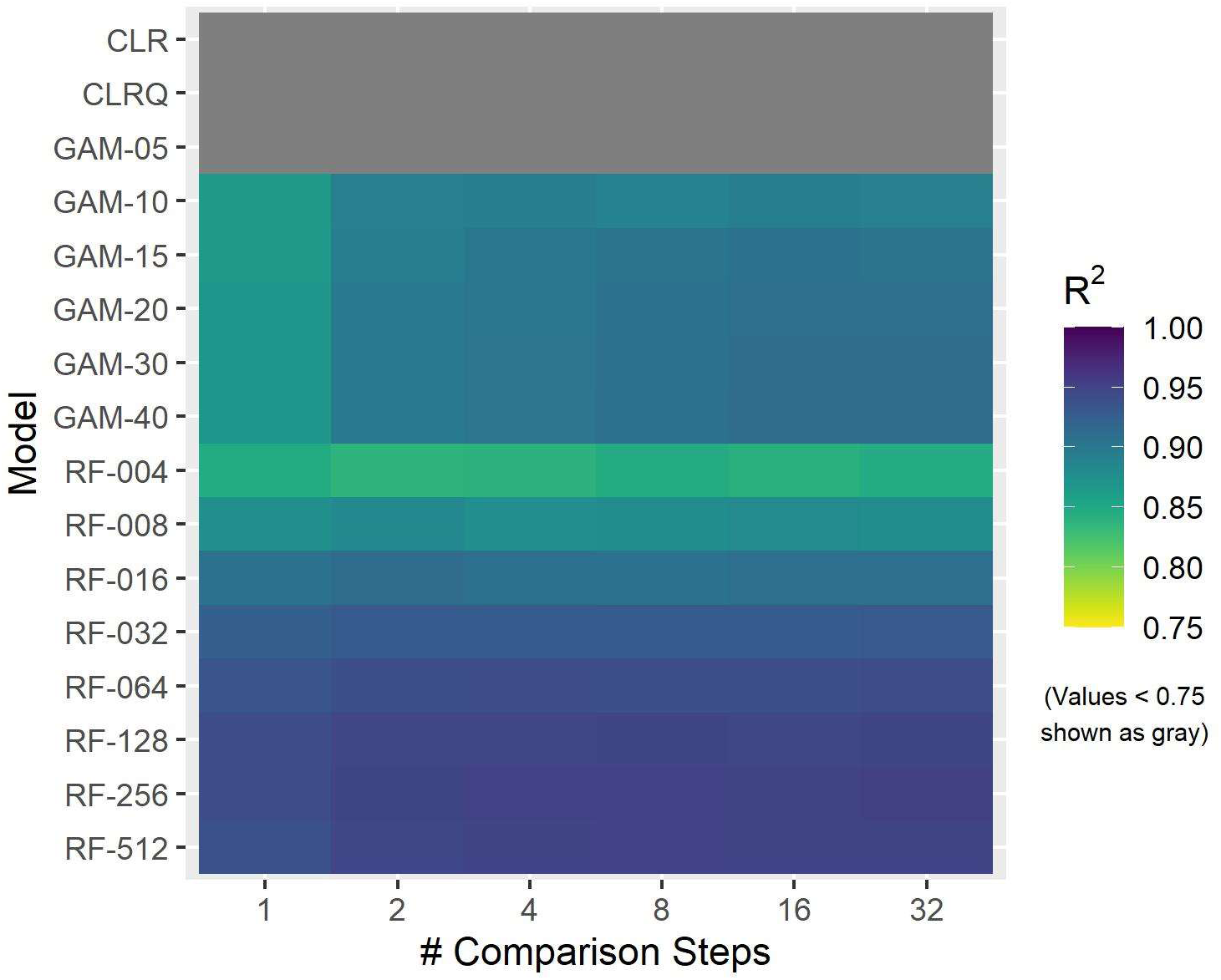
Heatmap of Mean Tuning-Set R^2^ Values for Different Models and Numbers of Comparison Steps. GAMs are labeled as *GAM-VW* and RFs as *RF-XYZ*, with *VW* indicating each spline’s degrees of freedom and *XYZ* indicating the number of OSs used to train each tree. To highlight differences between better values, only R^2^ values between 0.75 and 1 are depicted; values below 0.75 are shown in gray. Darker colors indicate better performance.

GAMs also generally perform best with larger numbers of CSs per OS. The best GAM has *k* = 40 dfs for each covariate’s spline (GAM-40) and 16 CSs per OS (mean R^2^ = 0.912, mean MPS = 0.112). As shown in Fig. S-6, while mean R^2^ generally increases with additional CSs, mean R^2^ appears to converge to similar values with 8 or more CSs for both RFs and GAMs; mean MPS shows similar trends (Fig. S-8).

CLR and CLRQ models showed less variation with different numbers of CSs, with R^2^ ranging from 0.228 to 0.246 regardless of the CSs. The best CLR model (R^2^ = 0.236, mean MPS = 0.265) has quadratic terms (i.e., CLRQ) and 32 CSs per OS.

Figure 4 gives examples of the effect curves yielded by these models. We show the effect curves for the tracks with step-down effects for which the RF-512 (8 CSs) model has its worst and best values of mean R^2^ minus MPS for (0.954 and 0.996 respectively for mean R^2^ and 0.016 and 0.000 for mean MPS). This difference (mean R^2^ minus MPS) characterizes the tradeoff between overall fit (R^2^) and general resemblance to the specified effect curve shape (MPS). These plots also contain effect curves from GAM-40 (16 CSs) and CLRQ models (32 CSs). The two plots show similar RF-based effect curves, reflecting the method’s consistency across the 25 tracks with this effect type. Compared with the other models’ effect curves, the RFs’ effect curves precisely capture the sharp drop in effect when the covariate value is 0.5. We show comparable plots for the other effect types in the Supplemental Material.

**Figure 4:**
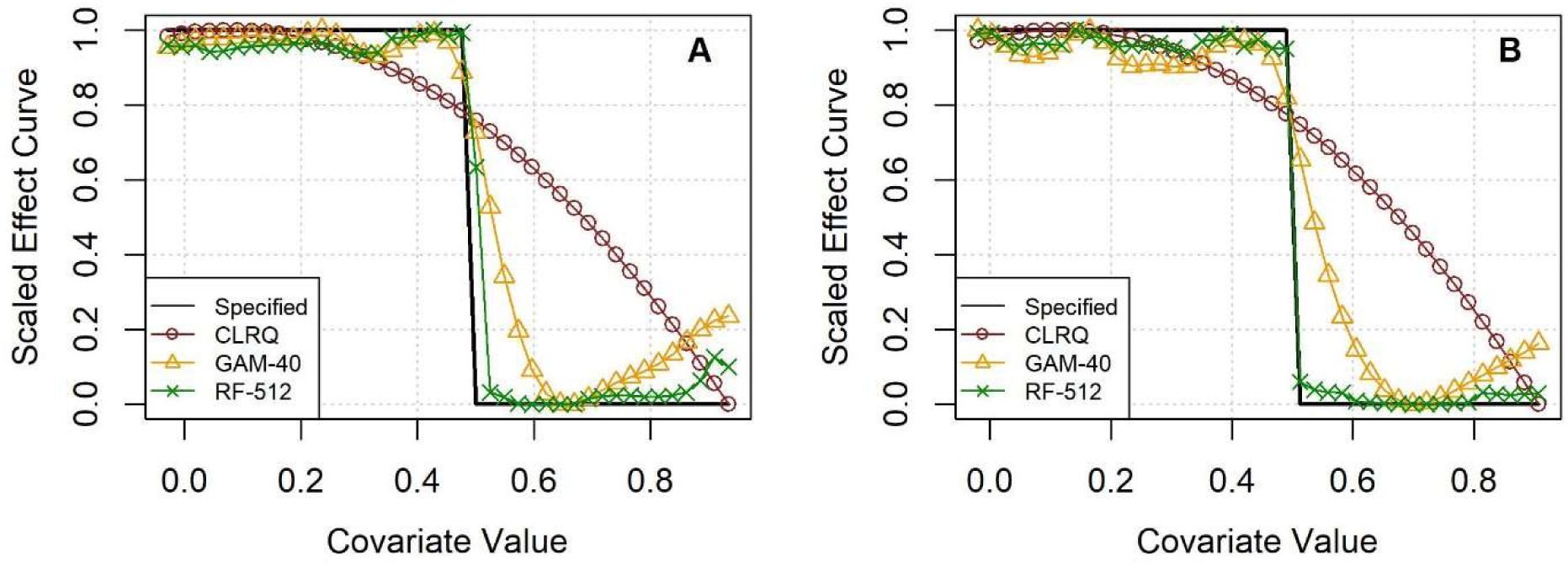
Effect Curve Examples for Tracks with Step-Down Effects with Best Model’s (A) Worst and (B) Best Performance.

Based on these results, we focus on the models from each type with the best settings. Figure 5 shows the results for each remaining model across all 100 tracks in the test set and for the 25 tracks with each of the effect types. To highlight smaller-scale differences, R^2^ values less than zero are not shown. Notably, larger values are better for R^2^, while smaller values are better for MPS. The RFs’ effect curves achieve the best results for all effect types except for linear effects.

**Figure 5:**
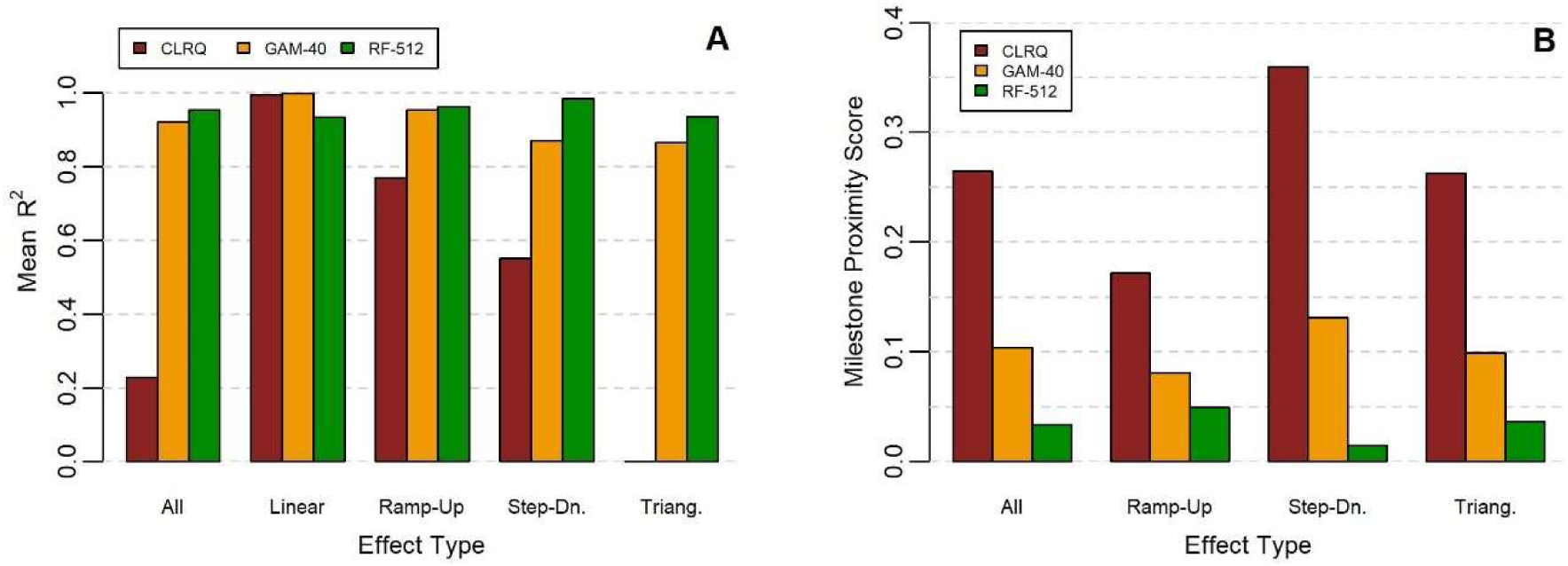
Test-Set Performance Comparison for Best Models of Each Type in Terms of (A) Mean R^2^ and (B) Mean Milestone Proximity Score. Larger values are better for R^2^. Smaller values are better for MPS.

Both for each effect type and overall, we use paired Wilcoxon signed rank (WSR) tests (Wilcoxon, 1992) to compare results from different pairs of models. For each of four effect types, there are 25 data points (from 5 landscapes and 5 tracks), resulting in 100 data points overall. We opted for the WSR tests because they do not require that the differences between models be normally distributed. We used two-sided tests to check for significant differences in either direction.

Table 1 shows the results of WSR tests. To conservatively account for testing multiple hypotheses simultaneously, we used 0.05 divided by the number of overall comparisons (18) as the threshold for significance (0.05/18 = 0.0028) (Peres-Neto, 1999). With 14 of 18 total comparisons, RF-512 is significantly better than the other models (10^-15^ < p < 0.001). In 2 comparisons (vs. GAM-40 for ramp-up effects), there are no statistically significant differences (0.012 < p ≤ 1.000). The CLRQ model and GAM-40 are significantly better than RF-512 in R^2^ for linear effects (10^-8^ < p < 0.0001); this is expected because the CLRQ model can be directly fitted to linear effects by setting the coefficients of the quadratic terms as zero. Specific p-values and mean differences from this analysis are included in Tables S-7 and S-8.

**Table 1:** Statistical Comparisons of Random Forests with Other Models.

| | $R^2$ | | Milestone Proximity Score | |
| --- | --- | --- | --- | --- |
|  | CLRQ | GAM-40 | CLRQ | GAM-40 |
| All | + | + | + | + |
| Linear | - | - | NA | NA |
| Ramp-Up | + | O | + | O |
| Step-Down | + | + | + | + |
| Triangular | + | + | + | + |
| “+”: RF-512 significantly better, “O”: no significant difference, “-”: RF-512 significantly worse, NA: no comparison possible |  |  |  |  |

More results from our simulation analysis, including heatmaps of results for each effect type, are reported in the Supplemental Material.

Figure 6 shows the results of applying our approach to wolf data (Barry et al., 2020). We constructed an RF-512 model, randomly sampling steps from across the 36 wolf tracks with 8 CSs paired with each OS.

**Figure 6:**
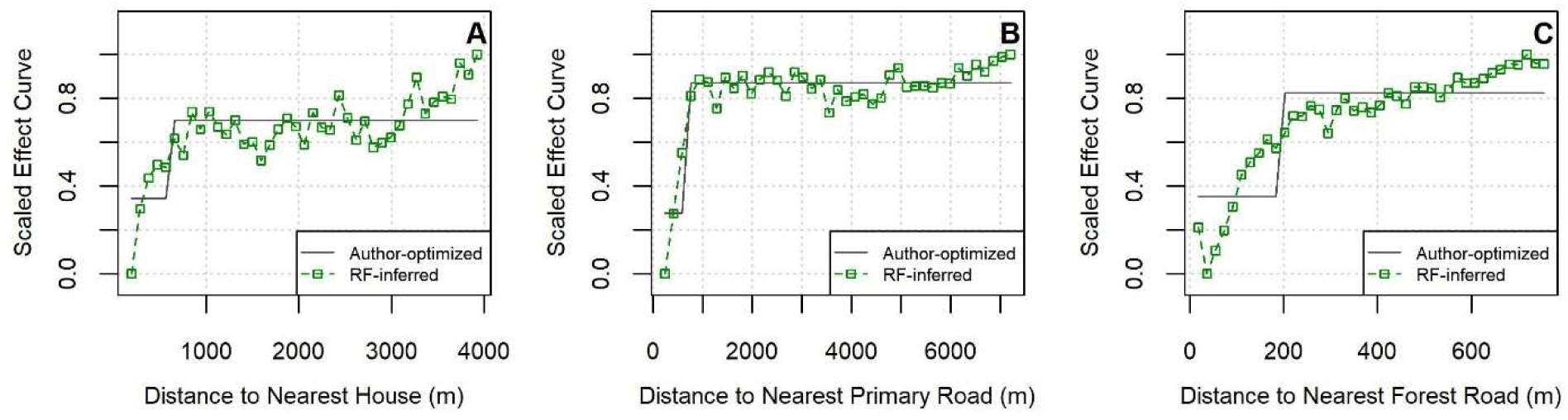
Effect Curves Inferred for Step Selection Covariates for Wolf Tracks, including (A) Distance to Nearest House, (B) Distance to Nearest Primary Road, (C) Distance to Nearest Forest Road. Data are from Barry et al. (2020).

The effect curves inferred by this RF resemble the step functions assumed by Barry et al. (2020) and seem to generally match their optimized thresholds. For illustration, we determined the values of the horizontal “steps” as the mean values of fitted effect curves before and after their optimized thresholds. Our models’ effect curves also confirm that the covariates have an approximate step-up effect (implying avoidance of houses and roads) rather than a step-down effect (which would imply attraction). At the same time, these effect curves illustrate the potential benefit of a RF approach, revealing that the effects of the analyzed distance covariates may follow a ramp function more closely than a step function.

Whether a step or ramp function, the RFs yield effect curves that stray somewhat from the flat step that would occur with larger covariate values (e.g., at 4000 m for distance to nearest house). Given the finite tracks and likelihood of unmodeled covariates also affecting wolves’ movement decisions, this variation is not surprising. Beyond the thresholds optimized by Barry et al. (2020), the RFs’ effect curves generally display the plateau that the authors had assumed, offering validation for the use of RFs for inferring nonlinear step selection effects.

## 4. Discussion

We have used simulated and real animal movement tracks to demonstrate the potential benefits of using RFs to analyze step selection. Compared with linear models, RFs offer the advantage of inference of nonlinear effects without the need to specify particular parametric forms. Moreover, our simulation analysis suggests that RFs significantly outperform CLR models with quadratic terms for ramp, step, and triangular covariate effects and outperform GAMs for step and triangular covariate effects. We also were able to roughly reproduce the findings of Barry et al. (2020) without invoking any subject matter expertise of the effects’ likely shapes.

This RF-based approach to SSFs has several possible uses. If nonlinear effects are suspected but not known, RFs offer a promising option for assessing particular relationships between covariate values and their effect on step selection. If a certain nonlinear effect type is hypothesized, RFs can likewise offer evidence for or against the hypothesis and also inform parameter decisions.

Using RFs to infer effect curves for step selection covariates is straightforward and can utilize available software. Given a track with OSs and corresponding CSs, the RF can be fit with the *randomForest* function within the *randomForest* package. While we used customized code to carefully compare our results with specified effect curves, effect curves for each covariate can similarly be computed with the *randomForest::partialPlot* function (Liaw & Wiener, 2002).

As described above, we found that the RF-512 (with 8 CSs) model yielded the best mean R^2^ and MPS across simulated tracks with all effect types. We selected this model based on tuning data and confirmed that it outperformed CLR models and GAMs on a separate set of test data. We also successfully replicated results from the real wolf tracks of Barry et al. (2020), using RFs to infer effect curves with shapes that closely resembled the step-up functions that Barry et al. (2020) had found.

We suspect that this particular RF-512 (with 8 CSs) achieved strong performance because of several characteristics: ample data to infer step selection trends; balance of OSs and CSs within the data used to train individual trees; and limited correlation between trees due to having a larger population of steps from which each tree’s training data could be sampled (made possible by having multiple CSs for each OS). Further study is needed to ascertain whether this model’s settings (8 CSs paired with each OS, evenly stratified sampling with replacement, 512 OSs and CSs to train each tree) also yield the best results on other data sets.

With our simulations, the number of OSs used to train each tree in the best RF (RF-512) equaled the number of OSs in each track. Thus, rather than using a fixed value (e.g., 512), an alternative strategy might involve setting the number of steps to train each tree as a given track’s number of strata (although this may pose a computational burden for longer tracks, e.g., with the wolf data, and introduce overfitting; compare Figs. S-20 and S-21).

In general, RFs have reasonable runtime, requiring more time than CLR models to train and evaluate, but considerably less time than GAMs (see Tables S-9 and S-10). Such efficiency is particularly important when working with longer movement tracks (which are increasingly available) or when fitting separate models for numerous animals as part of comparative research (which is increasingly popular).

There may be some effects for which RFs are not the best option. For instance, if effects are suspected to be linear, CLR models will likely yield the best results. Within our results, we found that CLR models yielded better performance on linear effects (as expected). Likewise, models with quadratic terms and GAMs yield smooth effect curves and may outperform RFs on inferring smoother effects (e.g., effects with higher-order parabolic curves). Nevertheless, our findings suggest that RFs offer a capable general-purpose approach for inferring effect curves when the shapes of the effects are unknown. Against each of the four types of effect that we studied, our best RF achieved a mean R^2^ of at least 0.933 (see Figure 5). In comparison, the GAM-40 and CLRQ models had mean R^2^ less than 0.871 for step-down and triangular effects.

Our promising results suggest that additional study with a wider range of tracks would be valuable. We used a limited range of simulation settings (exponentially distributed step length with mean = 4, von Mises concentration parameter= 3 for RTAs, track length = 512, smoothing window = 1, uncorrelated covariates, 3 total covariates). All of these settings likely affect model results. Thus, further work is needed to study these RF-based SSF options on additional simulated data. Likewise, for real wolf tracks, the RFs yielded effect curves that align wit hprior biological findings, but applying RFs to additional real track is warranted.

One limitation of our RFs is that there is no immediate process for determining the statistical significance of covariates within RFs. However, complementing our approach with a variable selection method like Boruta analysis (Kursa & Rudnicki, 2010) may provide comparable conclusions. Moreover, there are well-established permutation- and impurity-based metrics for assessing the “importance” of a covariate within a RF (Hastie et al., 2009).

Our approach also shares limitations with other step selection studies. Having data on the covariates that influence animals’ movement decisions is essential for accurate inference of covariate effects. No step selection analysis method can compensate for not having the covariates that drive movement decisions. Some factors that affect movement will inevitably omitted (e.g., due to challenges in measuring or collecting the values for such factors), but researchers should strive to include as many of the primary factors as possible.

It is also important to have tracks with sufficiently many OSs. Due to sampling noise, finite simulated tracks generally do not exactly reflect these specified effects. As long as tracks are “long enough,” major trends can generally be observed. However, as noise increases within an inferred effect curve, interpretation can be difficult.

RFs leverage the same data used to fit CLR-based SSFs: OSs complemented by CSs that feasibly could have been selected but were not observed. However, unlike CLR-based SSFs, RFs do not preserve the stratum structure of pairing individual OSs with their corresponding CSs. Accordingly, we also considered a novel variant of RFs in which the data for constructing each decision tree within a RF are adjusted to more closely resemble SSFs (see Section S-6). In particular, the data for training an individual trees would be drawn from one or more entire strata, rather than randomly from the entire set of available step data. The resulting “semi-matched random forest” (SMRF) models would mirror RFs in the use of random sampling (for strata for each tree and for covariates considered at each tree node) while still pairing observed and corresponding CSs (as with other SSFs). Ultimately, we found that SMRFs did not offer any advantage over RFs (see Section S-7), but we believe that the concept of preserving the stratum structure of the step data within the construction of RFs merits further exploration.

## 5. Conclusions

Two decades ago, Fortin et al. (2005) introduced the SSF framework of pairing OSs with CSs. Using this framework, dozens of studies have examined the factors that motivate animal movement decisions. Most of these studies have used CLR-based SSFs, which are constrained by the particular parametric form specified for each covariate; indeed, in many studies, only linear effects have been considered. Various enhancements have been proposed, including GAMs (Klappstein et al., 2024). Adding to this body of research, we have proposed RFs as an alternative to the traditionally used CLR-based SSFs. Using simulated and real data, we have demonstrated the potential advantages of this approach for inferring unknown nonlinear effects or assessing hypothesized nonlinear effects. Going forward, we envision the use of RFs as an approach that complements existing SSF approaches, providing insights about nonlinear relationships between covariate values and the effects that they have on animal movement decisions.

## Acknowledgments

PK and MR received funding from the Johns Hopkins Applied Physics Laboratory Innovation Program.

Author contributions

PK conceptualized the study and designed its methodology. PK and MR contributed to software and validation. PK curated the data and wrote the manuscript. All authors reviewed and edited the manuscript. WF provided supervision.

## Data availability

Simulated data can be obtained with the provided code. Data from the wolf case study are available at Mendeley (https://doi.org/10.17632/9722chrknz.1).

## Supplemental Material

### S-1. Overview of Analysis Approach

Figure S-1 provides a flowchart of how the components of our analysis are related to each other.

**Figure S-1:**
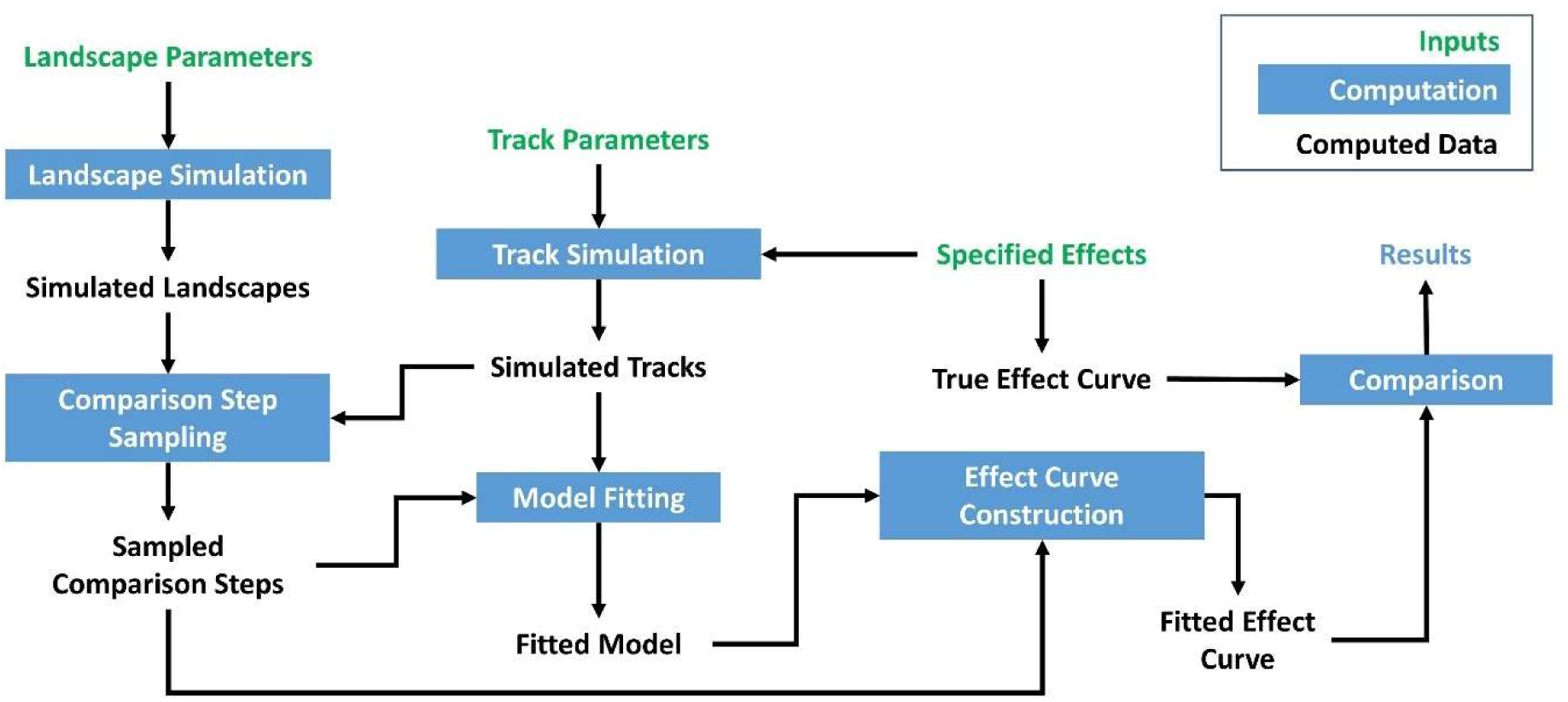
Flowchart of Analysis Approach.

### S-2. Additional Details on Simulations

We simulate one *G*-by-*G* grid of square cells for each covariate. To simulate an entire track without reaching the landscape boundary, we use *G =* 2001. For each grid cell, we sample the initial value from a uniform distribution from -1 to 2, which encompasses the different types of effects that we studied (see Table S-1). To reflect likely spatial correlation within the grid, we use a smoothing window with width 3, replacing each location’s original value with the mean of the original values within the 3-by-3 grid centered at the location. For cells along the boundary of the grid, we only consider the cells within a smoothing window that lie within the grid. Figure S-2 shows the values of such a grid, before and after smoothing.

**Figure S-2:**
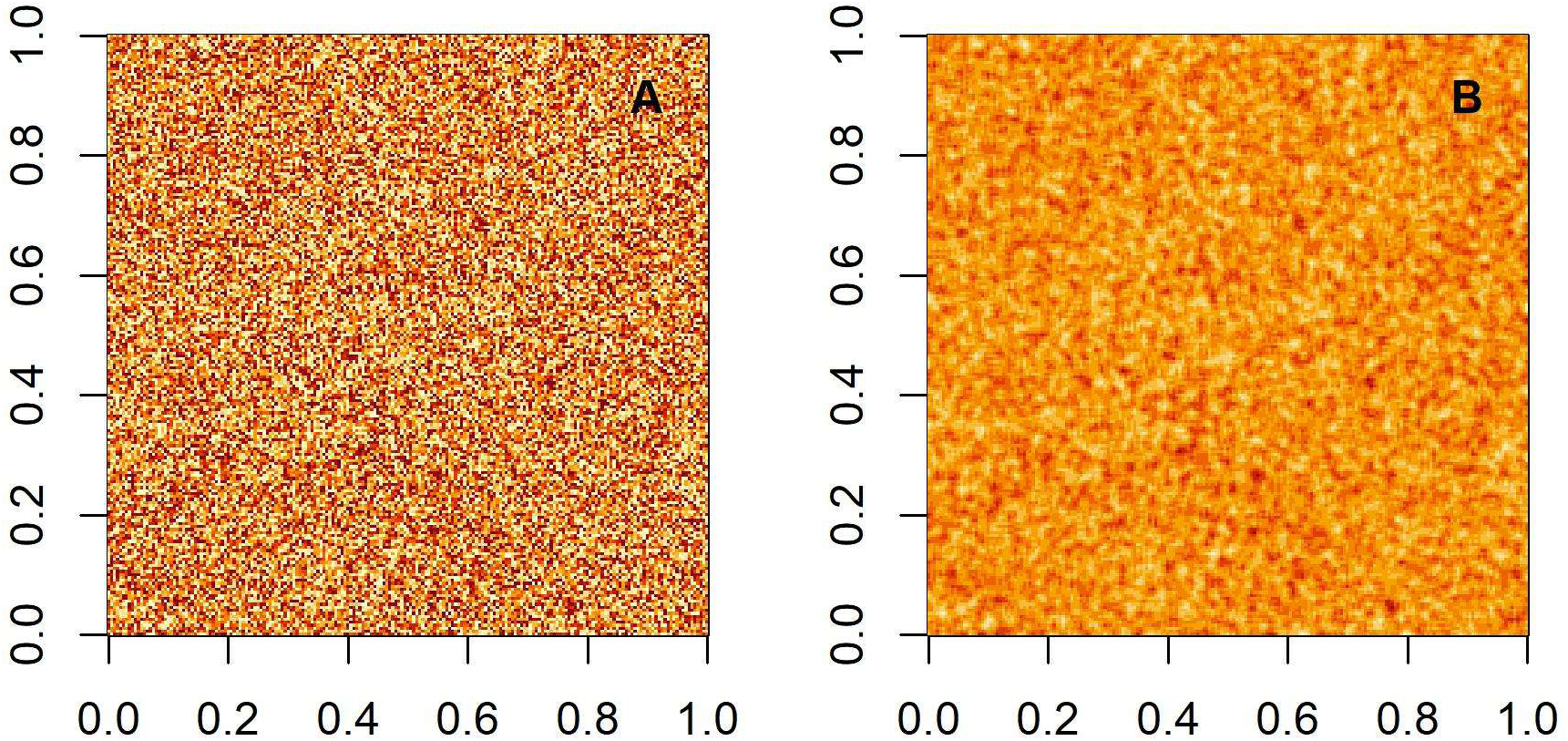
Example of Simulated Covariate Landscape with (A) Raw and (B) Smoothed Values.

This smoothing operation results in approximately normal distributions for each covariate (Agresti, 2018; Zhang et al., 2023). For illustration, see Figure S-3, which shows the distribution of values corresponding to Figure S-2 before and after smoothing.

**Figure S-3:**
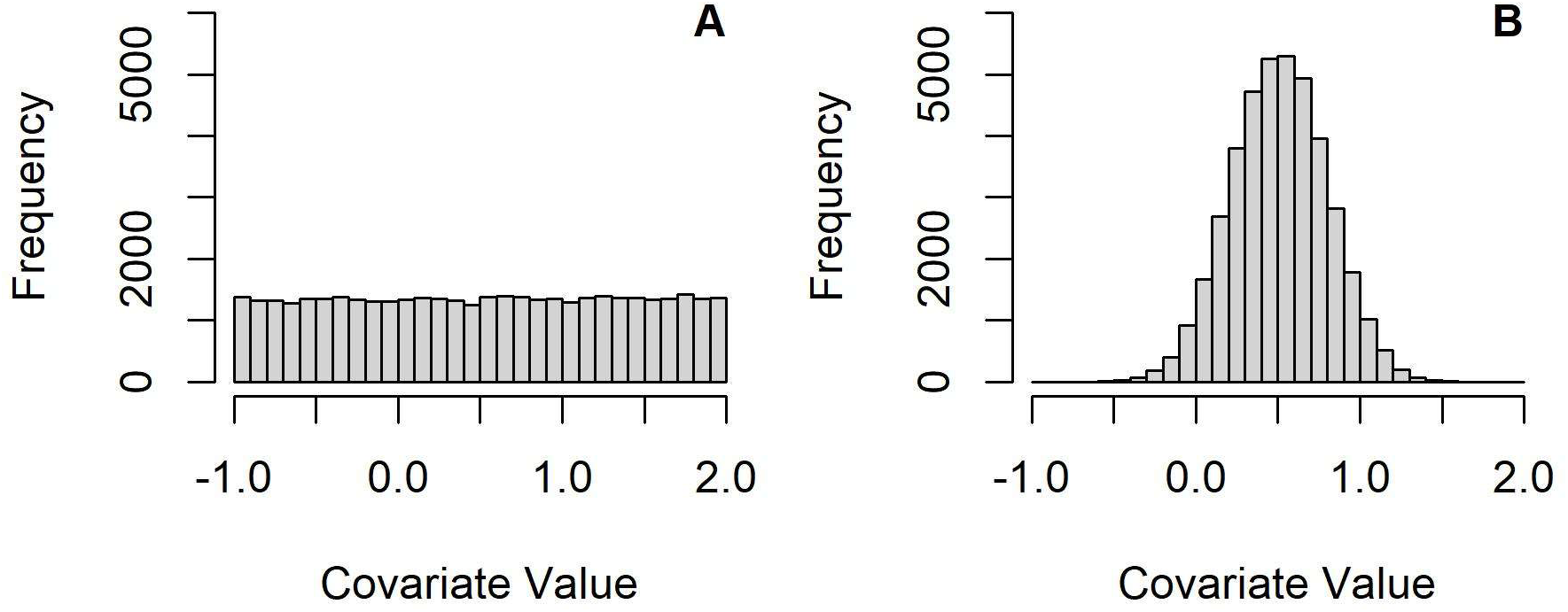
Histogram of (A) Raw and (B) Smoothed Simulated Covariate Values.

Given a simulated landscape of multiple covariates, we simulate tracks using the effect curves described in Table S-1 with the covariate values ***x*** for a given step contributing to the selection score *y*(***x***) for that step. We set various milestone points for each nonlinear effect type. Please see Section 2.5 for further details on computing the corresponding milestone proximity score (MPS).

**Table S-1:** Details for Each Effect Type.

| Effect Type | Equation for Contribution (y) to Selection Score for Covariate Value (x) | Milestone Points for Fitted and Scaled Effect Curve Values ($\hat{y}(x)$) with margin A |
| --- | --- | --- |
| Linear | $y = 4 * x$ | NA |
| Ramp-Up | $y = 0,$ if $x < 0.3$<br>$y = -3 + 10 * x,$ if $0.3 \leq x < 0.7$<br>$y = 4,$ if $x \geq 0.7$ | • Turn-up = $\max_{\hat{y}(x) \leq A} x$<br>Note: if $\hat{y}(x)$ is never below A, turn-up = min x<br>• Turn-flat = $\min_{\hat{y}(x) \geq 1 - A} x$<br>Note: if $\hat{y}(x)$ is never above $1 - A$, turn-flat = max x<br>• Ramp distance = turn-flat – turn-up |
| Step-Down | $y = 4,$ if $x < 0.5$<br>$y = 0,$ if $x \geq 0.5$ | • Step start = $\max_{\hat{y}(x) \geq 1 - A} x$<br>Note: if $\hat{y}(x)$ is never above $1 - A$, turn-up point = min x<br>• Step end = $\min_{x > \text{step start} \ \& \ \hat{y}(x) \leq A} x$<br>Note: if $\hat{y}(x)$ is never below A after the step start, step end = max x<br>• Step distance = step end – step start |
| Triangular | $y = 0,$ if $x < 0.3$<br>$y = -6 + 20 * x,$ if $0.3 \leq x < 0.5$<br>$y = 14 - 20 * x,$ if $0.5 \leq x < 0.7$<br>$y = 0,$ if $x \geq 0.7$ | • Max point = $mean \left( \max_{\hat{y}(x)=1} x \right)$<br>• Turn-up = $\max_{x < \text{max point} \ \& \ \hat{y}(x) \leq A} x$<br>Note: if $\hat{y}(x)$ is never below A prior to max point, step end = min x<br>• Flat-out = $\min_{x > \text{max point} \ \& \ \hat{y}(x) \leq A} x$<br>Note: if $\hat{y}(x)$ is never below A after max point, step end = max x<br>• Triangle distance = flat-out – turn-up |

Please see Figure S-4 for an example of an individual track. The track from a particular simulation is determined by the underlying covariate landscapes and the random sampling of the candidate steps.

**Figure S-4:**
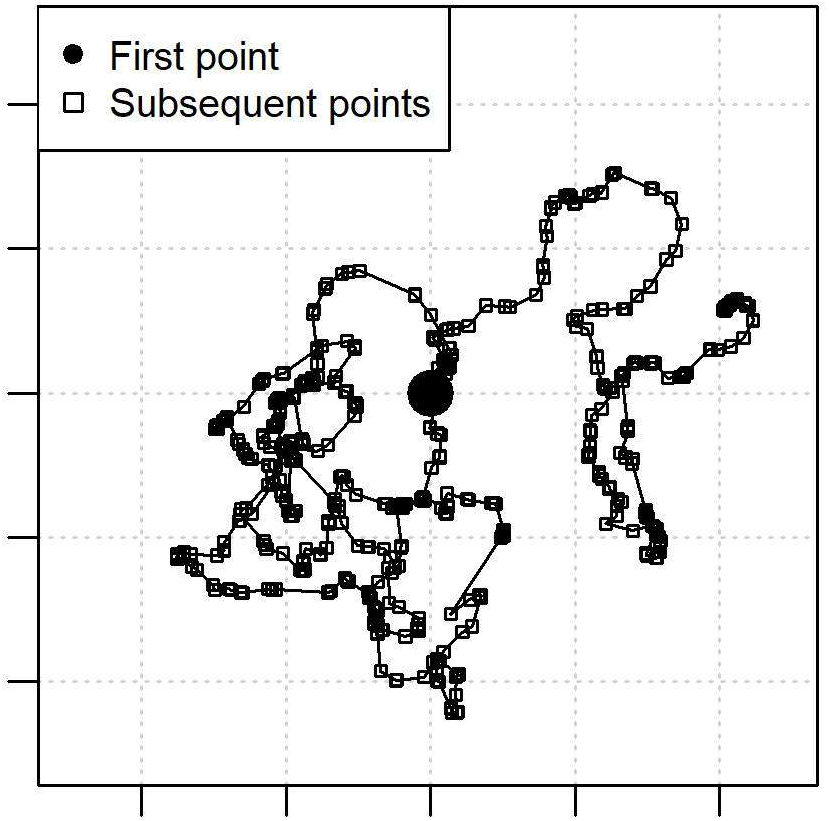
Example of a Simulated Track.

### S-3. Review of Random Forests

Training a random forest (RF) model consists of the following steps:

1. Gather training data, e.g., strata of comparison steps (CSs) paired with each observed step (OS)
2. Set parameters (number of data points to sample for each tree, sampling scheme, number of covariates to consider at each split, number of training set examples that determine when a branch has reached a “terminal node”)
3. For each tree, draw a sample from all training data
4. At the first split, sample some number of candidate covariates
5. For each covariate under consideration, sort the values from the data for training that tree (i.e., sampled in step 3); find values that lie between each pair of consecutively ordered values; assess how well these values split the data
6. Find the best value (among all values from all covariates under consideration) at splitting the data
7. Repeat for all splits in the tree, continuing until all branches have terminal nodes
8. Repeat for all trees in the forest

Notably, the process for constructing a semi-matched random forest (SMRF) (Section 4 and Section S-6) is identical to this process for constructing a RF with the exception of step 3. For SMRFs, the data sampled in step 3 for each tree come from one or more strata.

Each stratum of steps consists of an OS paired with one or more CSs.

### S-4. Sensitivity Analysis for Model Parameters

We conduct sensitivity analysis on several decisions related to our models’ setup: (A) Sampling scheme for RFs; (B) the value of the smoothing constant for RFs and GAMs.

#### S-4.1 Sampling Scheme for Random Forests

Given *M* CSs per OS, we consider three options for determining the step type distribution for the steps used to train each tree:

1. no stratification and randomly sampling 1 + *M* steps;
2. representative stratification with 1 OS and *M* CSs sampled for each tree; and
3. even stratification with 1 OS and 1 CS sampled for each tree.

Notably, the amount of step data from which the training data are sampled increases with *M* for all three options, but the amount of data used to train each tree only increases with *M* for the first two options.

We also assess sampling with and without replacement for the training steps for each tree. When steps are sampled with replacement, a given step may be sampled more than once for a given tree. Such repeated steps reduce the amount of unique data used to train each tree, but also increase the decorrelation between trees.

Table S-2 and Table S-3 summarizes the mean R^2^ for models run with *M* = 8 CSs on the full set of 100 tracks. The values corresponding to the selected settings are shown in boldface.

**Table S-2:** Mean R^2^ with Different Training Data Sampling Schemes for Random Forest Trees.

| Step Type Distribution per Tree | No Stratification (1 + $M$ CSs sampled randomly) | Representative Stratification (All CSs + 1 observed step) | Even Stratification (1 CS + 1 observed step) |
| --- | --- | --- | --- |
| Without Replacement | 0.646 | 0.646 | 0.926 |
| With Replacement | 0.657 | 0.655 | **0.936** |

**Table S-3:** Mean Milestone Proximity Score with Different Training Data Sampling Schemes for Random Forest Trees.

| Step Type Distribution per Tree | No Stratification (1 + $M$ CSs sampled randomly) | Representative Stratification (All CSs + 1 observed step) | Even Stratification (1 CS + 1 observed step) |
| --- | --- | --- | --- |
| Without Replacement | 0.0830 | 0.0837 | 0.0572 |
| With Replacement | 0.0802 | 0.0784 | **0.0568** |

With even stratification, sampling with replacement is significantly better in R^2^ than sampling without replacement (p < 10^-5^) according to the one-sample Wilcoxon signed rank (WSR) test. There is no significant difference in MPS (p = 0.81) between these options.

Based on these results, we opt to use even stratification and replacement to sample the steps for each tree within our random forest models (Section 2.3).

#### S-4.2 Smoothing Constant for Computing Effect Curves

The logarithmic function is undefined for zero or infinity. Therefore, to transform the output score *μ* of a given model with the half-logit function 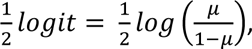 we need to replace values of *μ* that are close to 0 or 1. We achieve this by selecting a smoothing constant *C* such that all values of *μ* less than *C* are replaced by *C* and all values of *μ* greater than 1-*C* are replaced by 1-*C* (see Section 2.4). This smoothing step ultimately yields step functions with plateaus for output scores near 0 or 1. The width of the plateau is determined by *C*; larger values of *C* have wider plateaus. Moreover, smaller values of *C* permit output scores near 0 or 1 to be “stretched” father away from the other transformed values. Figure S-5 illustrates this phenomenon. In this figure, *eps* refers to 2.2*10^-16^.

**Figure S-5:**
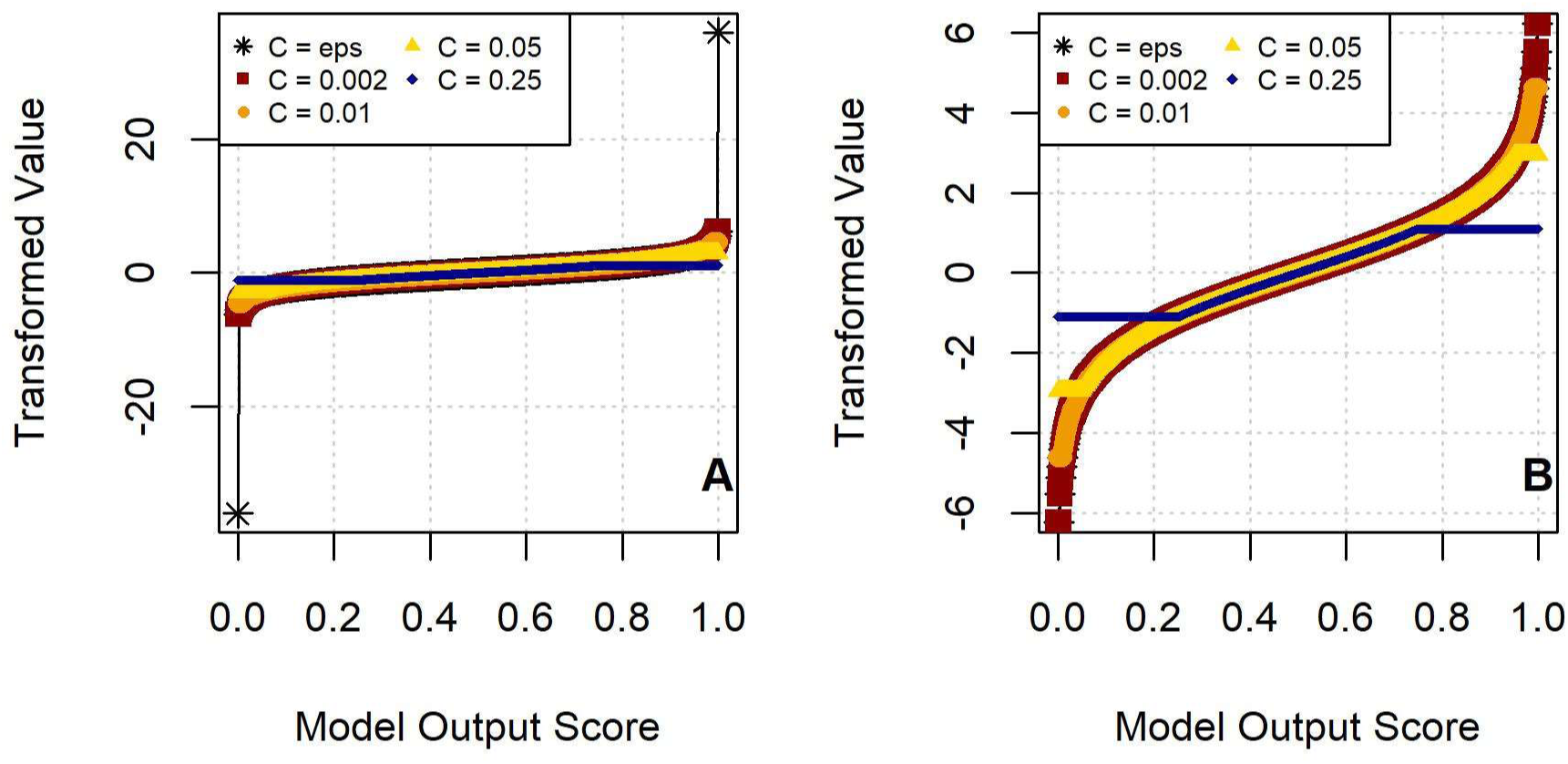
Effect on Half-Logit Transformation of Different Smoothing Constants, (A) Full Scale and (B) Zoomed In.

We assess the effect of different candidate values for *C*: 2.2*10^-16^, 0.002, 0.01, 0.05, 0.25. Output scores for RFs are discrete, reflecting the number of trees that have “voted” for the positive class. For RFs with 500 trees, *C* = 0.002 amounts to assigning the same transformed values for steps with votes from 0 or 1 trees. Likewise, *C* = 0.01 amounts to all vote totals from 0 to 5 trees having the same transformed value. Output scores of generalized additive models (GAMs) and conditional logistic regression (CLR) models are continuous-valued and can fall anywhere within 0 and 1.

We consider the effect of *C* on both RFs and GAMs. For RFs, we used *M* = 8 CSs, evenly stratified sampling with replacement (see Section S-4.1), and 512 OSs and 512 CSs per tree. For GAMs, we use *M* = 8 CSs and *k* = 15 degrees of freedom. Table S-4 shows the results from this analysis. The best results for each model type are shown in boldface. (See also Section 2.4.)

**Table S-4:** Mean R^2^ with Different Smoothing Constants.

| Smoothing Constant $C$ | RF | GAM |
| --- | --- | --- |
| $2.2 \times 10^{-16}$ | 0.936 | **0.902** |
| 0.002 | 0.936 | 0.893 |
| 0.01 | 0.947 | 0.888 |
| 0.05 | **0.951** | 0.881 |
| 0.25 | 0.814 | 0.881 |

Based on these results, we opt to use *C =* 0.05 for RFs and *C =* 2.2*10^-16^ (*.Machine$double.eps* in R) for GAMs.

### S-5. Additional Details for Constructing Effect Curves

Constructing an effect curve for a given covariate consists of the following steps:

1. Identify a covariate of interest
2. Find the 5% and 95% quantiles among all observed values for that covariate (including OSs and CSs)
3. Find *N_EC_* evenly spaced values spanning the interval bounded by these quantiles (we used *N_EC_* = 40)
4. Find all combinations of other features from all steps (OSs and CSs) from all strata
5. Pair all covariate-of-interest values with all combinations of other features, resulting in (1 + *M*) * (# strata) combinations with each value of the covariate of interest
6. For a given value of the covariate of interest, run all combinations with that value through the model, tracking the model’s output score (e.g., the proportion of trees that predicted the observed-step class) for each combination
7. Compute the mean output score *μ* for that value of the covariate of interest
8. If *μ* < *C*, replace the *μ* with *C*; if *μ* > 1 – *C*, replace *μ* with 1 – *C*
9. Apply the following transformation: 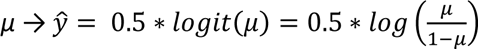
10. Repeat for all *N_EC_* values of the covariate of interest
11. If comparing effect curves from multiple models, scaled all transformed values with 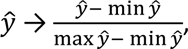 thus projecting all values to [0, 1]

Step 5 involves pairing all values of the covariate of interest with all combinations of all other covariates. This may yield combinations that never actually occur in practice. With certain covariates, some resulting combinations may be very improbable (e.g., high elevation and low distance to water).

One alternative approach is to pair a given value of the covariate of interest only with the combinations of other covariates that coincide with covariate-of-interest values that are close to the given value. However, this limits the number of combinations for each covariate-of-interest value and may lead to uneven numbers of combinations for different covariate-of-interest values. Criteria for what determines when two values are “close” must also be set. Moreover, the observed combinations of values may not be representative of the full population of combinations that could occur.

Another alternative approach is to fit a joint probability distribution for all covariates. With the value of the covariate of interest fixed, the values of the other covariates would be drawn from the resulting conditional distribution. This approach also has drawbacks, though, as fitting a joint distribution is not always straightforward. Not all combinations of covariates may be compatible with a specific parametric form, e.g., multivariate normal. Nonparametric approaches, e.g., kernel density, likewise may not be representative of the full population of combinations that could occur.

### S-6. Semi-Matched Random Forests

RFs leverage the same data used to fit step selection function (SSFs) based on CLR: OSs complemented by CSs that feasibly could have been selected but were not observed. However, unlike SSFs, they do not preserve the stratum structure that keeps together steps with a common starting point, i.e., individual OSs and their corresponding CSs. Accordingly, we also considered a novel variant of RFs in which the data for constructing each decision tree within a RF are adjusted to more closely resemble SSFs.

In particular, the data for training an individual trees would be drawn from one or more entire strata, rather than randomly from the entire set of available step data. The resulting “semi-matched random forest” (SMRF) models would mirror RFs in the use of random sampling (for strata for each tree and for covariates considered at each tree node) while still pairing OSs and CSs (as with other SSFs).

For SMRFs, we construct one tree at a time, providing only the step data from the sampled strata to fit that tree. We consider sampling 1, 2, 4, 8, 16, 32, 64, 128, 256, and 512 strata to train each tree, sampling these strata randomly and with replacement. We refer to such models as SMRF-*XYZ*-R, where *XYZ* indicates the number of sampled strata. “R” indicates that the strata are sampled randomly, rather than requiring that the strata be consecutive within the track.

To conveniently include the other features of RFs (random sampling of covariates at each split), we use the *randomForest* function in the *randomForest* package (Liaw & Wiener, 2002) to construct each tree as a one-tree RF (setting *ntree* to 1) before aggregating the trees into a single 500-tree RF. Unlike RFs, we use all steps from a stratum to train a given tree within a SMRF (enforced by setting the *sampsize* in the *randomForest* function to the number of steps within that stratum). Like RFs, we randomly sample which covariates to consider at each tree split.

While some SMRFs outperform CLR models and GAMs (Table S-5), they offer no improvement over RFs. The benefit of retaining such a “step starting point in common” is less than we had anticipated. Thus, rather than SMRFs, we have focused this paper on applying well-established RFs to SSFs. Nevertheless, we believe that the concept of preserving the stratum structure of the step data within the construction of SMRFs has conceptual appeal and merits further exploration.

**Table S-5:** Mean R^2^ for All Effect Types on Tuning Set.

| Model \ #CSs | 1 | 2 | 4 | 8 | 16 | 32 |
| --- | --- | --- | --- | --- | --- | --- |
| CLR | 0.242 | 0.241 | 0.243 | *0.245* | 0.244 | 0.242 |
| CLRQ | 0.228 | 0.236 | 0.235 | 0.233 | 0.234 | **0.236** |
| GAM-05 | 0.637 | 0.647 | 0.640 | 0.624 | 0.630 | 0.598 |
| GAM-10 | 0.866 | 0.892 | 0.893 | 0.890 | 0.893 | 0.892 |
| GAM-15 | 0.866 | 0.896 | 0.900 | 0.904 | 0.906 | 0.905 |
| GAM-20 | 0.868 | 0.898 | 0.902 | 0.907 | 0.910 | 0.910 |
| GAM-30 | 0.869 | 0.898 | 0.903 | 0.908 | 0.912 | 0.912 |
| GAM-40 | 0.869 | 0.898 | 0.903 | 0.908 | **0.912** | 0.912 |
| RF-004 | 0.847 | 0.838 | 0.840 | 0.847 | 0.841 | 0.846 |
| RF-008 | 0.875 | 0.882 | 0.876 | 0.878 | 0.882 | 0.878 |
| RF-016 | 0.906 | 0.913 | 0.908 | 0.907 | 0.910 | 0.910 |
| RF-032 | 0.926 | 0.930 | 0.929 | 0.928 | 0.928 | 0.931 |
| RF-064 | 0.935 | 0.941 | 0.941 | 0.940 | 0.940 | 0.942 |
| RF-128 | 0.941 | 0.948 | 0.948 | 0.949 | 0.947 | 0.949 |
| RF-256 | 0.941 | 0.949 | 0.951 | 0.951 | 0.950 | *0.952* |
| RF-512 | 0.938 | 0.947 | 0.950 | **0.951** | 0.950 | 0.951 |
| SMRF-001 | 0.751 | 0.585 | 0.416 | 0.270 | 0.113 | -0.154 |
| SMRF-002-R | 0.806 | 0.712 | 0.528 | 0.431 | 0.178 | -0.157 |
| SMRF-004-R | 0.843 | 0.760 | 0.668 | 0.526 | 0.253 | -0.045 |
| SMRF-008-R | 0.876 | 0.841 | 0.772 | 0.641 | 0.390 | 0.109 |
| SMRF-016-R | 0.912 | 0.892 | 0.839 | 0.729 | 0.515 | 0.284 |
| SMRF-032-R | 0.927 | 0.912 | 0.869 | 0.765 | 0.575 | 0.349 |
| SMRF-064-R | 0.934 | 0.929 | 0.883 | 0.781 | 0.594 | 0.363 |
| SMRF-128-R | 0.940 | 0.935 | 0.891 | 0.784 | 0.599 | 0.399 |
| SMRF-256-R | *0.940* | **0.934** | 0.886 | 0.774 | 0.595 | 0.412 |
| SMRF-512 | 0.936 | 0.928 | 0.875 | 0.759 | 0.584 | 0.435 |

Unlike RFs, SMRFs seem to perform better with fewer CSs paired with each OS. We suspect that the improvement with fewer CSs may come from the increased class balance in the steps used to train each tree. When using entire strata to train a given tree, having fewer CSs is the only way to increase class balance.

### S-7. Additional Results

Table S-5 shows the values underlying Fig. 3, as well as values from SMRFs. The results corresponding to the settings with the highest mean R^2^ for each type of model are shown with italics. The values corresponding to the selected settings for each model type (determined by both R^2^ and MPS) are shown in boldface.

Figure S-6 shows the effect on mean R^2^ of different numbers of CSs for RF-512 and GAM-40 models, illustrating the limited benefit (for these particular simulated tracks) of having more than 8 CSs paired with each OS. Having 4 or 8 CSs yields mean R^2^ values that are nearly as high as with 16 or 32 CSs.

**Figure S-6:**
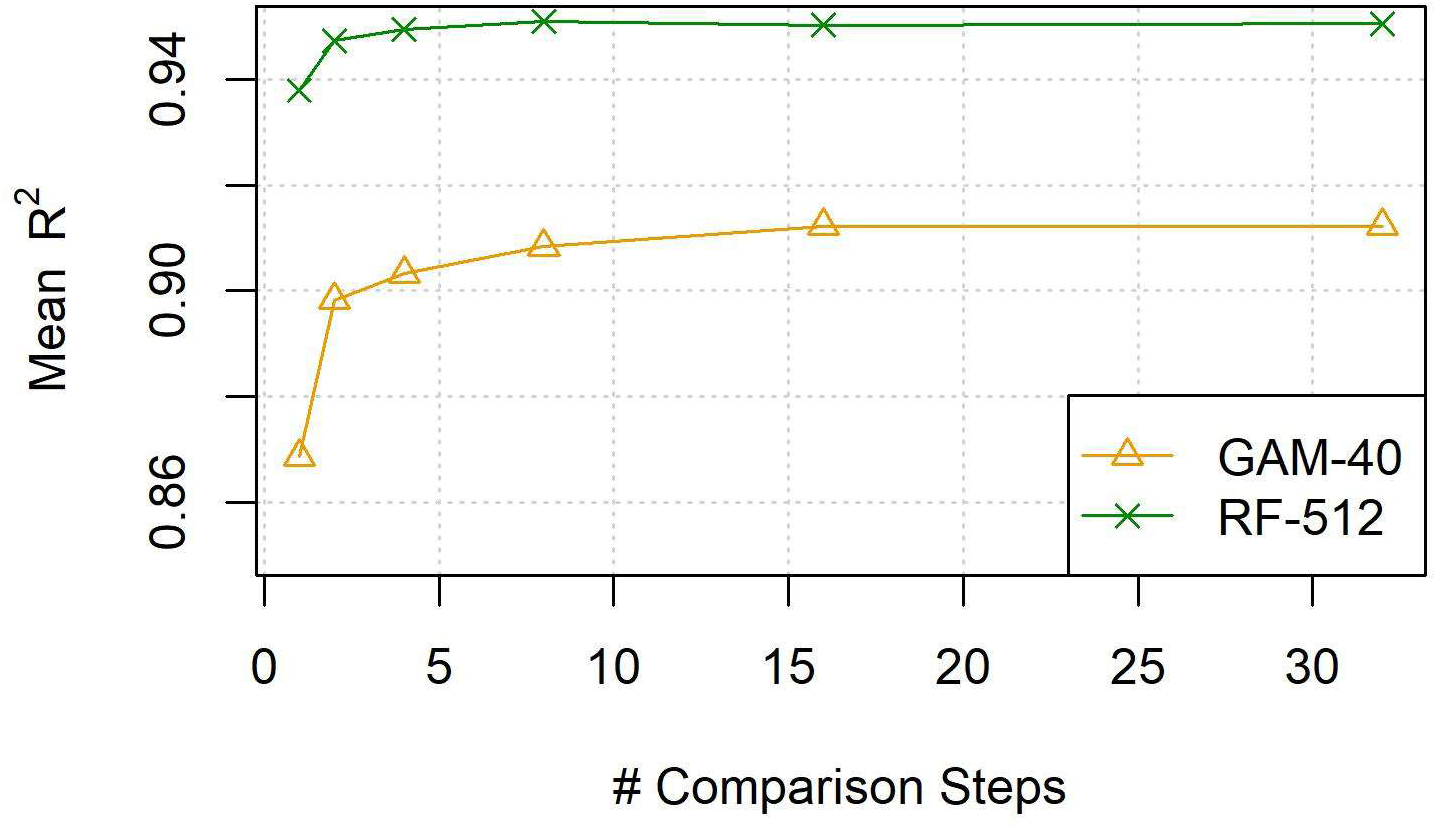
Mean R^2^ vs. Number of Comparison Steps for Best Performing Models.

Complementing the mean R^2^ values in Fig. 3, Figure S-7 shows the mean MPSs for different model types and numbers of CSs. Darker colors indicate better performance. RFs show the strongest performance with this metric.

**Figure S-7:**
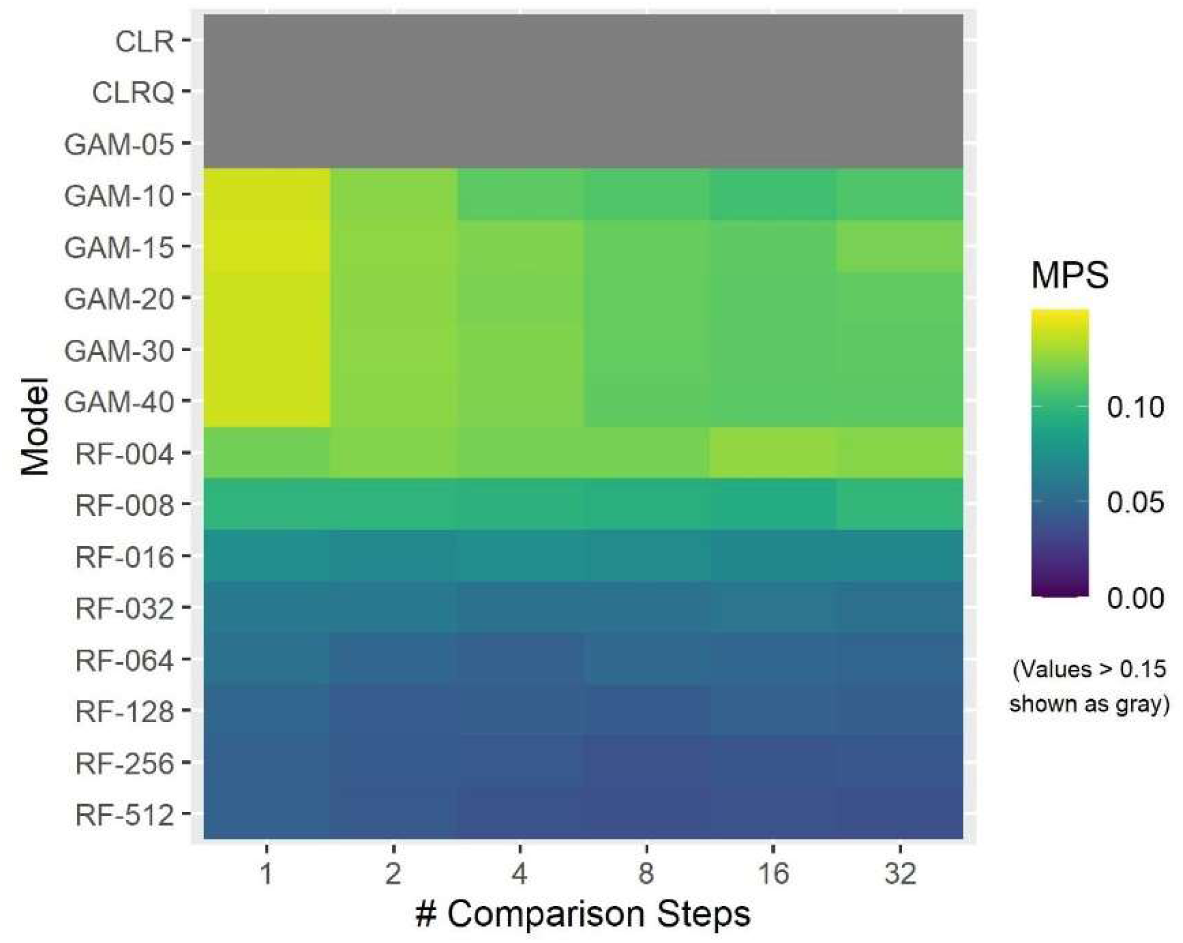
Heatmap of Mean Tuning-Set Milestone Proximity Scores for Different Models and Numbers of Comparison Steps.

Table S-6 shows the values underlying Figure S-7. The results corresponding to the settings with the highest mean MPS for each type of model are shown with italics. The values corresponding to the selected settings for each model type (determined by both R^2^ and MPS) are shown in boldface.

**Table S-6:** Mean Milestone Proximity Score for All Effect Types on Tuning Set.

| Model \ #CSs | 1 | 2 | 4 | 8 | 16 | 32 |
| --- | --- | --- | --- | --- | --- | --- |
| CLR | 0.390 | 0.388 | 0.389 | 0.389 | 0.389 | 0.389 |
| CLRQ | 0.276 | 0.271 | 0.268 | 0.266 | 0.265 | ***0.265*** |
| GAM-05 | 0.202 | 0.195 | 0.193 | 0.190 | 0.189 | 0.193 |
| GAM-10 | 0.140 | 0.124 | 0.113 | 0.110 | *0.105* | 0.109 |
| GAM-15 | 0.140 | 0.125 | 0.121 | 0.115 | 0.113 | 0.120 |
| GAM-20 | 0.138 | 0.124 | 0.120 | 0.115 | 0.112 | 0.114 |
| GAM-30 | 0.139 | 0.124 | 0.122 | 0.114 | 0.113 | 0.113 |
| GAM-40 | 0.139 | 0.124 | 0.121 | 0.113 | 0.112 | 0.112 |
| RF-004 | 0.118 | 0.122 | 0.119 | 0.119 | 0.126 | 0.123 |
| RF-008 | 0.098 | 0.097 | 0.097 | 0.095 | 0.092 | 0.099 |
| RF-016 | 0.075 | 0.071 | 0.074 | 0.072 | 0.070 | 0.070 |
| RF-032 | 0.060 | 0.060 | 0.057 | 0.056 | 0.058 | 0.055 |
| RF-064 | 0.057 | 0.049 | 0.046 | 0.051 | 0.049 | 0.048 |
| RF-128 | 0.048 | 0.044 | 0.045 | 0.043 | 0.046 | 0.045 |
| RF-256 | 0.046 | 0.042 | 0.041 | 0.038 | 0.040 | 0.040 |
| RF-512 | 0.046 | 0.041 | 0.038 | ***0.036*** | 0.038 | 0.036 |
| SMRF-001 | 0.150 | 0.195 | 0.202 | 0.204 | 0.215 | 0.226 |
| SMRF-002-R | 0.143 | 0.172 | 0.196 | 0.198 | 0.212 | 0.220 |
| SMRF-004-R | 0.120 | 0.153 | 0.173 | 0.194 | 0.206 | 0.213 |
| SMRF-008-R | 0.101 | 0.097 | 0.133 | 0.170 | 0.197 | 0.214 |
| SMRF-016-R | 0.073 | 0.061 | 0.073 | 0.109 | 0.158 | 0.184 |
| SMRF-032-R | 0.060 | 0.049 | 0.053 | 0.074 | 0.095 | 0.123 |
| SMRF-064-R | 0.053 | 0.043 | 0.048 | 0.068 | 0.086 | 0.104 |
| SMRF-128-R | 0.052 | 0.041 | 0.046 | 0.072 | 0.086 | 0.092 |
| SMRF-256-R | 0.047 | ***0.036*** | 0.047 | 0.071 | 0.080 | 0.090 |
| SMRF-512 | 0.046 | 0.038 | 0.052 | 0.070 | 0.086 | 0.088 |

Figure S-8 shows the effect on mean MPS of different numbers of CSs for RF-512 and GAM-40 models, again illustrating the limited benefit (for these particular simulated tracks) of having more than 8 CSs paired with each OS.

**Figure S-8:**
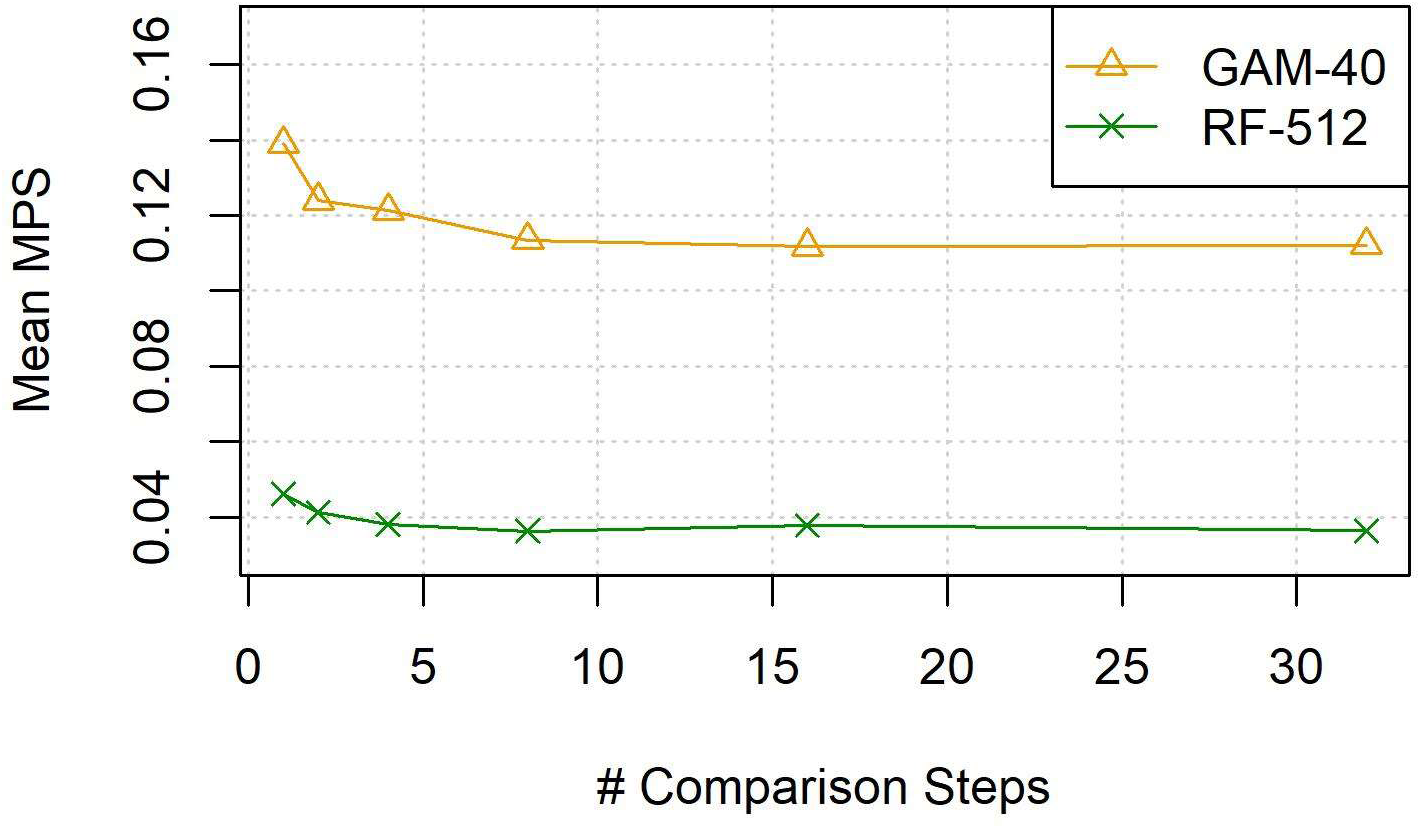
Mean Milestone Proximity Score vs. Number of Comparison Steps for Best Performing Models.

For linear effects (Figure S-9), CLR models and GAMs have strong performance. For instance, GAM-10 with 32 CSs has mean R^2^ = 0.998. Some RFs also perform fairly well; RF-256 and RF-512 with 2 or more CSs have mean R^2^ > 0.920. (Because MPS is designed to assess how well fitted effect curves resembled specified nonlinear effects, we did not calculate MPS for linear effects.)

**Figure S-9:**
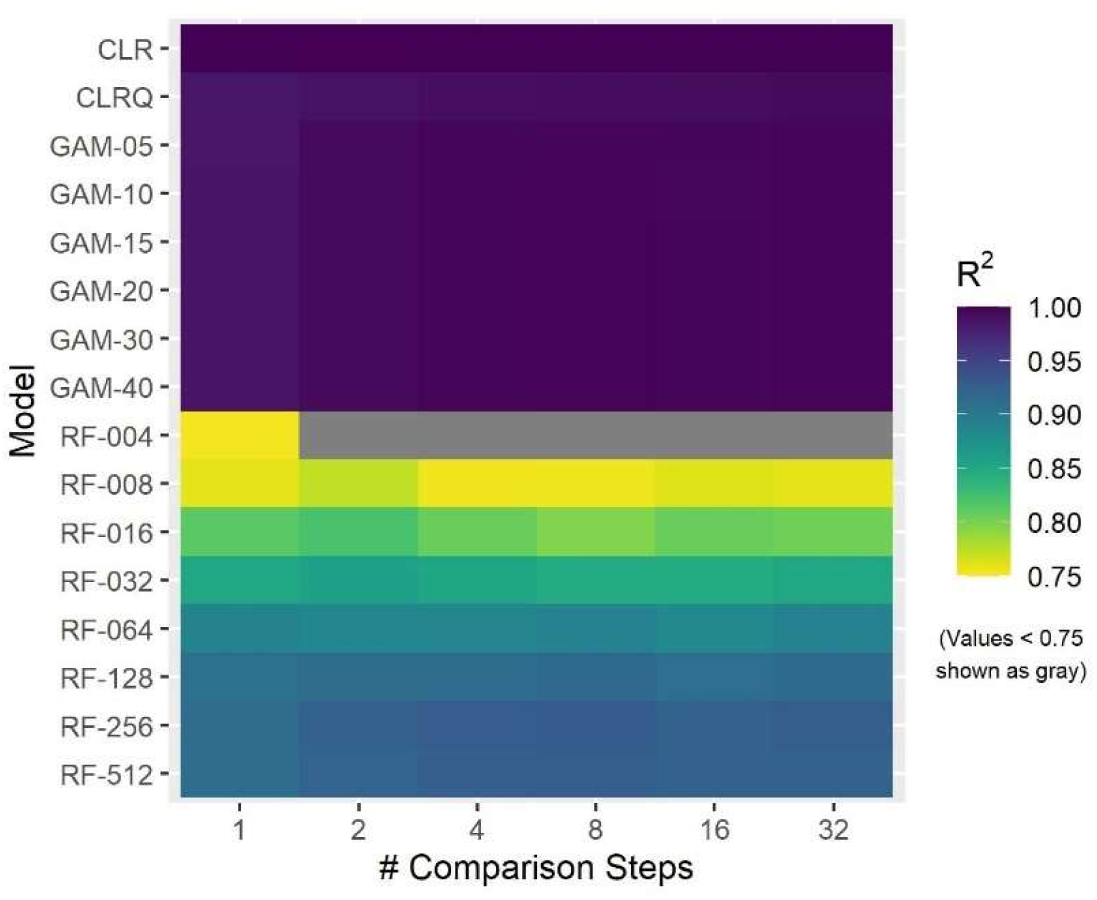
Heatmap of Mean Tuning-Set R^2^ Values for Different Models and Numbers of Comparison Steps for Tracks with Linear Effects.

Against ramp-up effects (Figure S-10 and Figure S-11), GAMs exhibit good performance (for instance, mean R^2^ = 0.961 for GAM-15 with 4 CSs), perhaps due to the resemblance of the ramp-up curve to a smooth shape. RFs show the best performance; RF-256 (32 CSs) has mean R^2^ = 0.964 and mean MPS = 0.046. CLR models have less strong performance as the effect deviates from a linear shape.

**Figure S-10:**
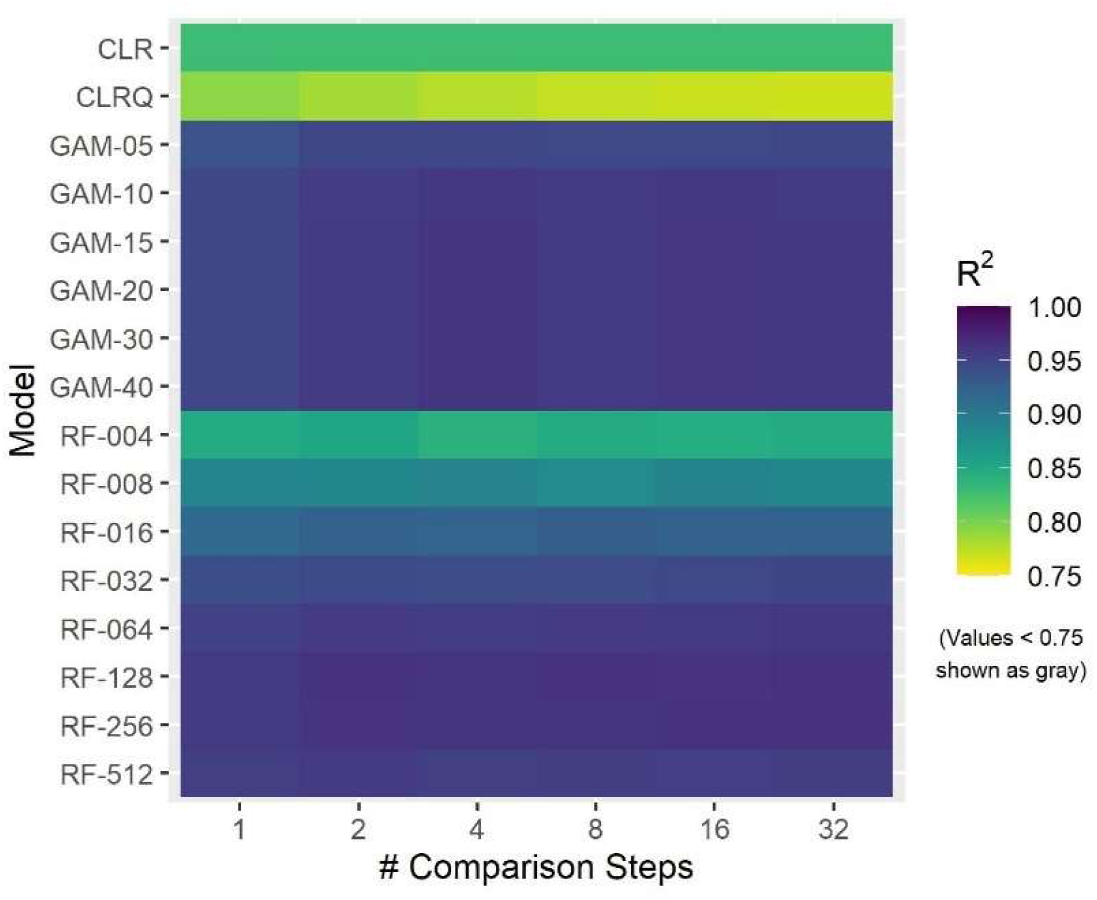
Heatmap of Mean Tuning-Set R^2^ Values for Different Models and Numbers of Comparison Steps for Tracks with Ramp-Up Effects.

**Figure S-11:**
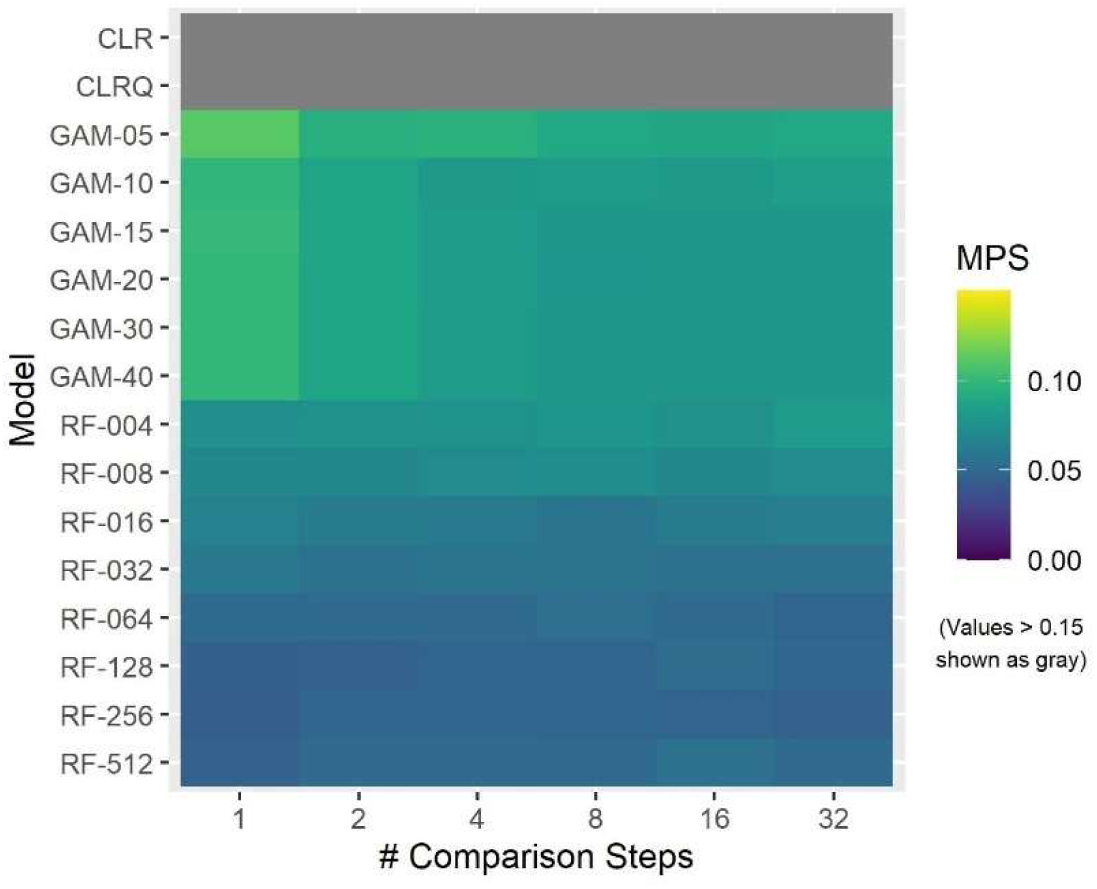
Heatmap of Mean Tuning-Set Milestone Proximity Scores for Different Models and Numbers of Comparison Steps for Tracks with Ramp-Up Effects.

With step-down effects (Figure S-12 and Figure S-13), RFs have the strongest performance. RF-512 (32 CSs) has mean R^2^ = 0.983 and mean MPS = 0.013. Mean R^2^ values for GAMs with *k* ≥ 10 range from 0.799 to 0.867 with the best performance achieved by GAM-40 with 32 CSs. CLR models show poorer performance with step-down effects with mean R^2^ ranging from 0.535 to 0.667.

**Figure S-12:**
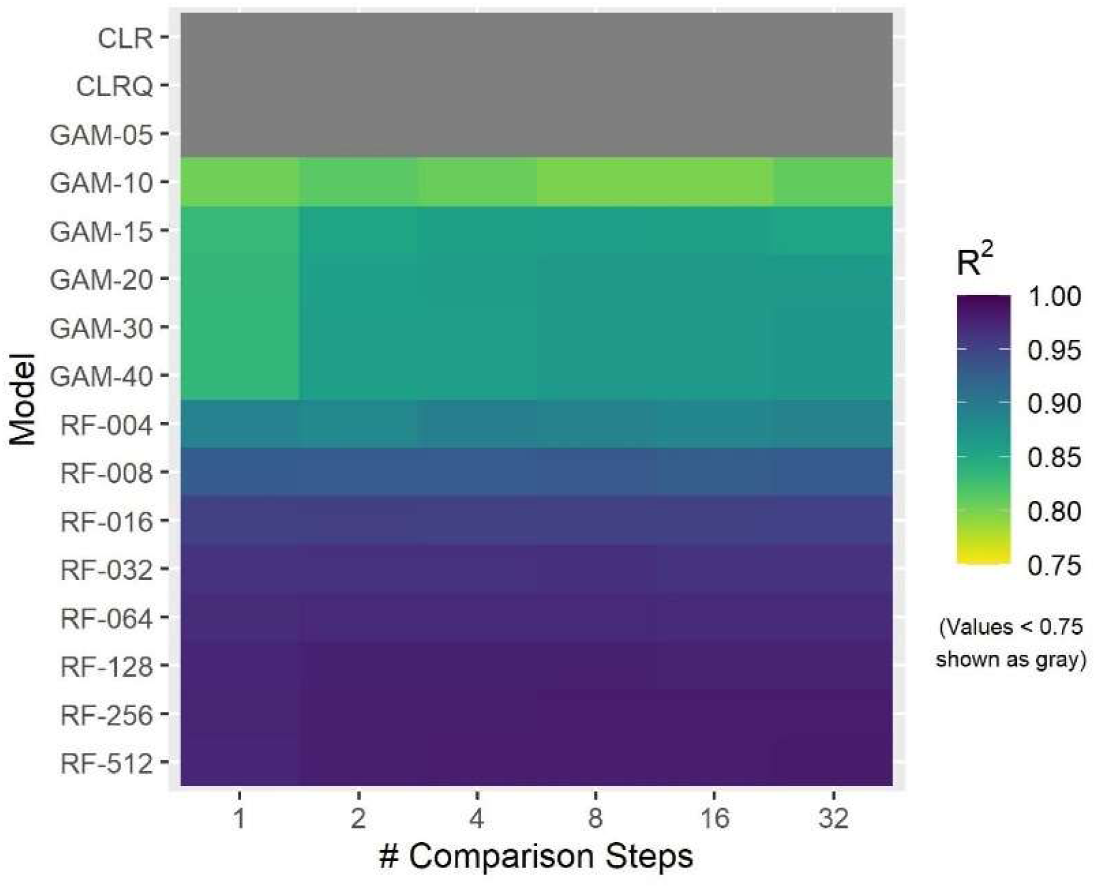
Heatmap of Mean Tuning-Set R^2^ Values for Different Models and Numbers of Comparison Steps for Tracks with Step-Down Effects.

**Figure S-13:**
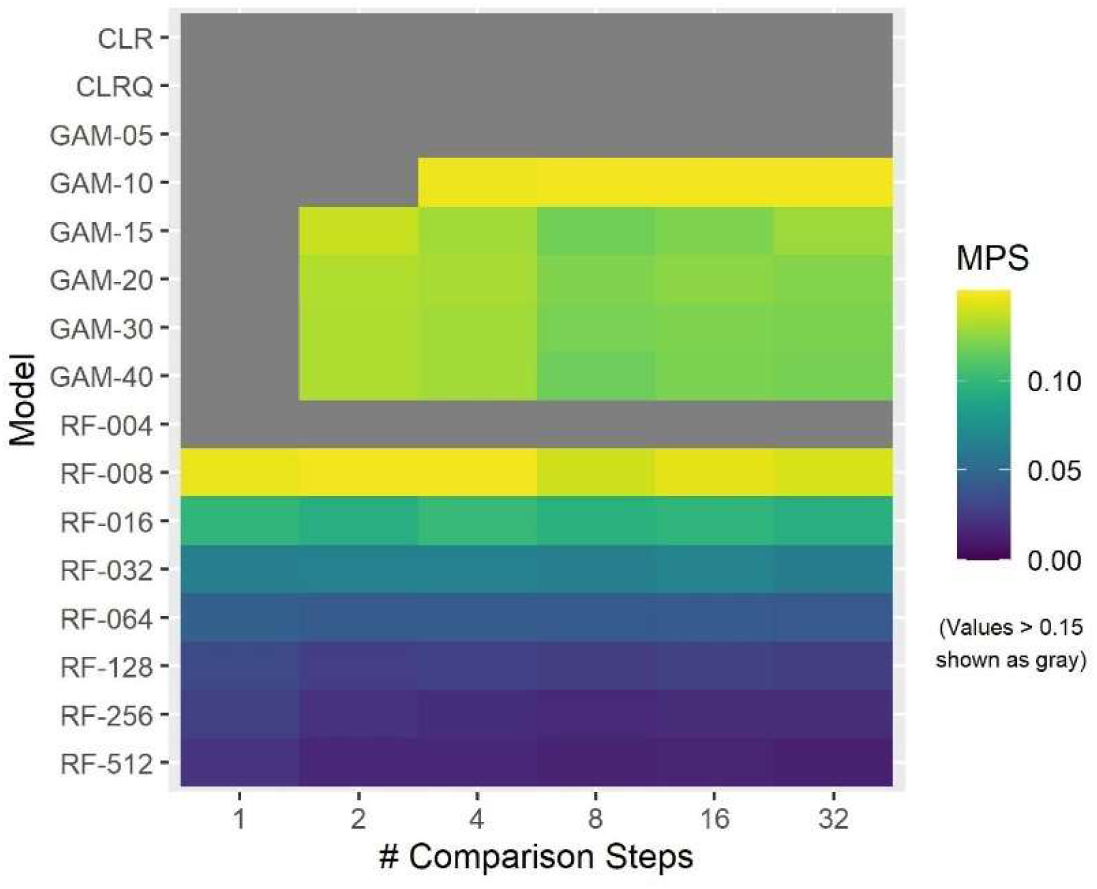
Heatmap of Tuning-Set Milestone Proximity Scores for Different Models and Numbers of Comparison Steps for Tracks with Step-Down Effects.

With triangular effects (Figure S-14 and Figure S-15), RFs again have the strongest performance. RF-032 with 4 CSs has mean R^2^ = 0.960 and mean MPS = 0.047. RF-256 with 8 CSs (which has the best performance across all effect types) has mean R^2^ = 0.940 and mean MPS = 0.040. GAMs seem to correctly infer the preference for intermediate values but miss the sharp change. For GAMs with *k* ≥ 10 and at least 4 CSs, mean R^2^ ranges from 0.784 to 0.826 and mean MPS from 0.086 to 0.153. CLR models have poor performance against triangular effects with mean R^2^ less than 0, indicating that their errors are higher than those of a model that merely predicted a constant value.

**Figure S-14:**
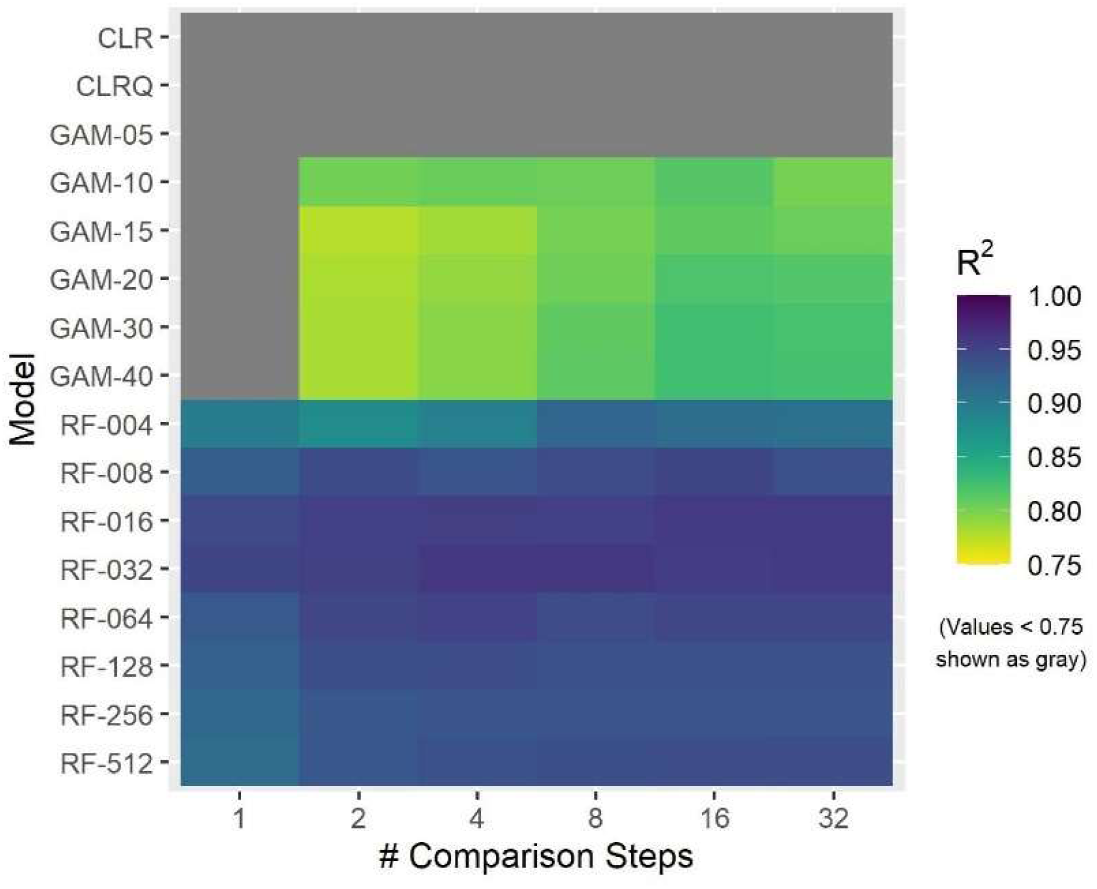
Heatmap of Mean Tuning-Set R^2^ Values for Different Models and Numbers of Comparison Steps for Tracks with Triangular Effects.

**Figure S-15:**
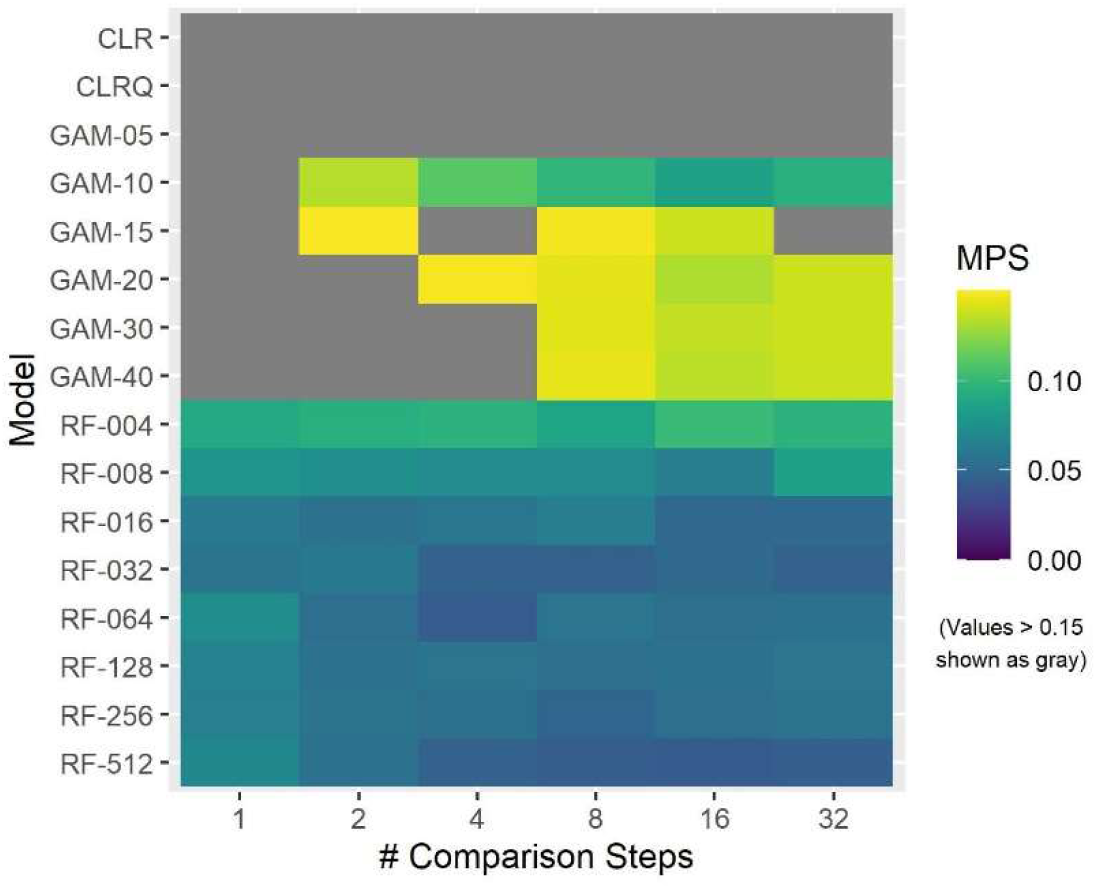
Heatmap of Mean Tuning-Set Milestone Proximity Scores for Different Models and Numbers of Comparison Steps for Tracks with Triangular Effects.

Figure S-16 through Figure S-18 provide examples of fitted effect curves from for linear, ramp-up, and triangular effects respectively. See Fig. 4 for examples of effect curves for step-down effects. Each figure shows the effect curves from the tracks of that effect type for which RF-512 (8 CSs) had its worst and best performance (in terms of R^2^ minus MPS). The figures also show effect curves from GAM-40 (16 CSs) and CLRQ (32 CSs) models.

**Figure S-16:**
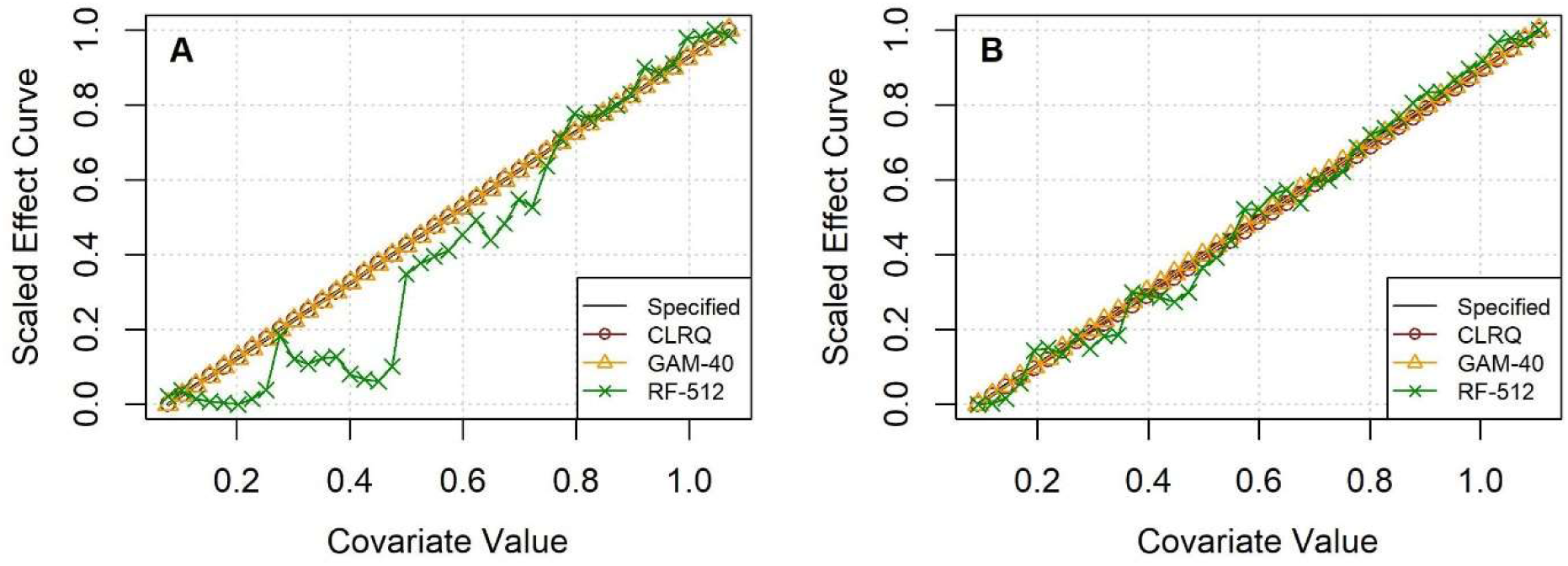
Effect Curve Examples for Tracks with Linear Effects with Best Model’s (A) Worst and (B) Best Performance.

**Figure S-17:**
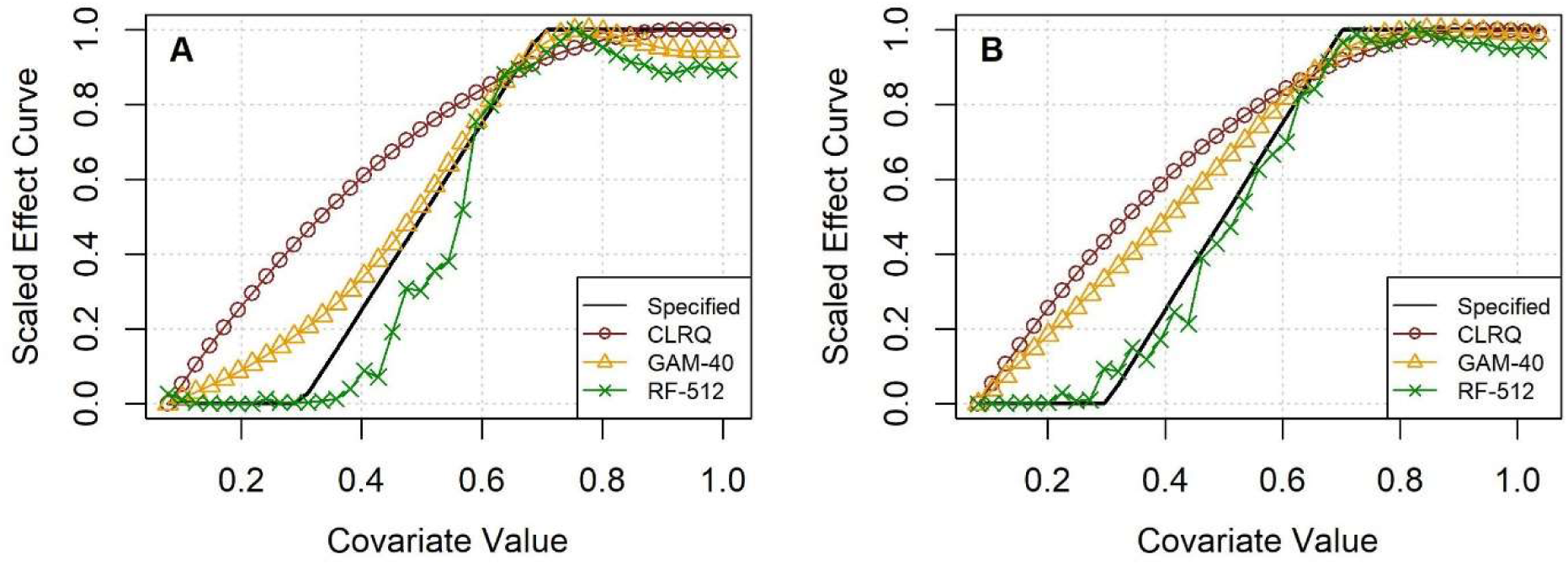
Effect Curve Examples for Tracks with Ramp-Up Effects with Best Model’s (A) Worst and (B) Best Performance.

**Figure S-18:**
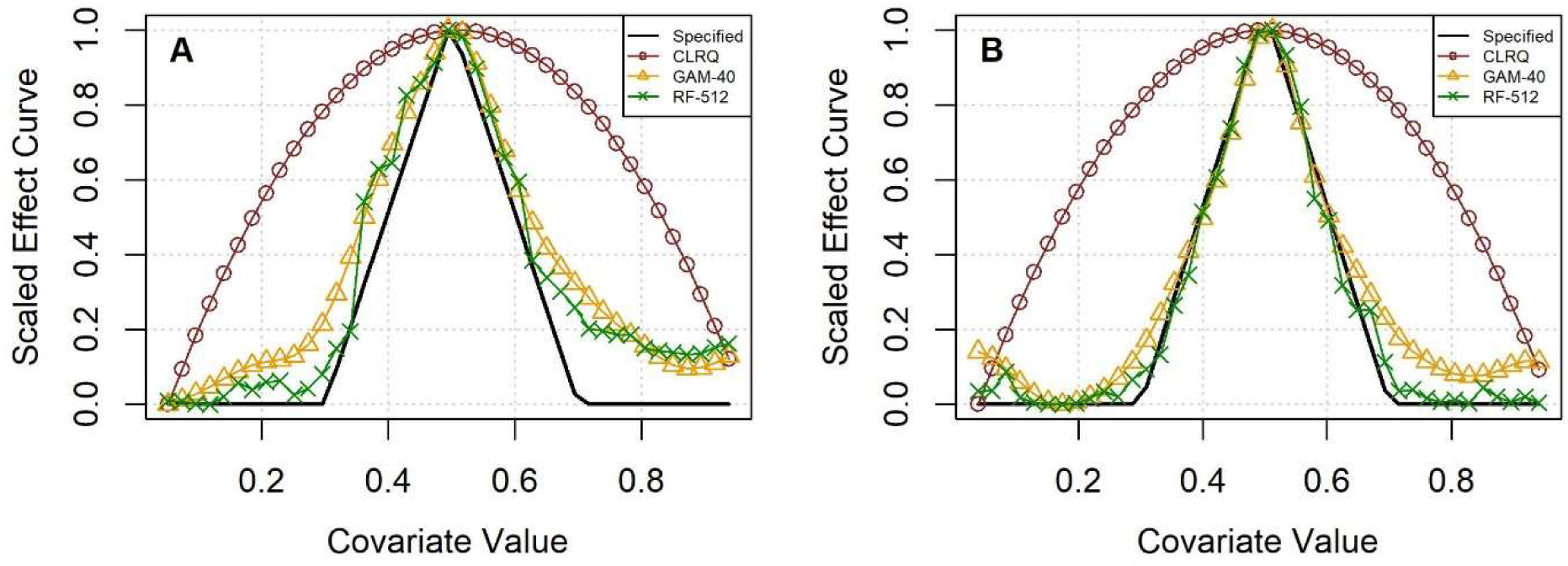
Effect Curve Examples for Tracks with Triangular Effects with Best Model’s (A) Worst and (B) Best Performance.

Table S-7 and Table S-8 give the p-values and mean differences corresponding to the statistical tests shown in Table 1.

**Table S-7:** P-Values from Wilcoxon Signed Rank Tests Comparing RF-512 with Other Models.

|  | R<sup>2</sup> |  | Milestone Proximity Score |  |
| --- | --- | --- | --- | --- |
|  | CLRQ | GAM-40 | CLRQ | GAM-40 |
| All | $4.5 * 10^{-14}$ | $9.6 * 10^{-4}$ | $5.3 * 10^{-14}$ | $8.1 * 10^{-11}$ |
| Linear | $6.0 * 10^{-8}$ | $1.3 * 10^{-5}$ | NA | NA |
| Ramp-Up | $6.0 * 10^{-8}$ | 1.00 | $1.2 * 10^{-5}$ | 0.0122 |
| Step-Down | $6.0 * 10^{-8}$ | $1.3 * 10^{-5}$ | $1.2 * 10^{-5}$ | $1.3 * 10^{-5}$ |
| Triangular | $6.0 * 10^{-8}$ | $5.2 * 10^{-5}$ | $1.3 * 10^{-5}$ | $9.7 * 10^{-5}$ |

**Table S-8:** Mean Differences in Performance Metrics (RF-512 minus Other Model)

|  | R<sup>2</sup> |  | Milestone Proximity Score |  |
| --- | --- | --- | --- | --- |
|  | CLRQ | GAM-40 | CLRQ | GAM-40 |
| All | 0.72 | 0.03 | -0.23 | -0.07 |
| Linear | -0.06 | -0.06 | NA | NA |
| Ramp-Up | 0.19 | 0.01 | -0.12 | -0.03 |
| Step-Down | 0.43 | 0.11 | -0.34 | -0.12 |
| Triangular | 2.34 | 0.07 | -0.23 | -0.06 |

Table S-9 and Table S-10 compare the times required to fit each type of model and construct the corresponding effect curves. The times corresponding to the best settings of each model type are shown in boldface. The times for the best RF are considerably faster than the times for the best GAM. These tables illustrate how runtimes increase with additional CSs and greater model complexity (with more degrees of freedom for GAMs and more data used to train each tree for RFs).

**Table S-9:** Mean Time to Fit Model (in seconds)

| Model \ #CSs | 1 | 2 | 4 | 8 | 16 | 32 |
| --- | --- | --- | --- | --- | --- | --- |
| CLR | 0.03 | 0.03 | 0.04 | 0.06 | 0.13 | 0.37 |
| CLRQ | 0.03 | 0.04 | 0.06 | 0.08 | 0.17 | **0.46** |
| GAM-05 | 1.97 | 1.94 | 2.35 | 3.40 | 17.30 | 66.76 |
| GAM-10 | 2.84 | 2.82 | 3.35 | 4.61 | 19.83 | 76.19 |
| GAM-15 | 4.78 | 4.61 | 5.19 | 6.30 | 25.42 | 96.97 |
| GAM-20 | 22.29 | 21.66 | 22.63 | 14.66 | 83.78 | 313.17 |
| GAM-30 | 51.07 | 54.80 | 64.56 | 28.05 | 109.60 | 442.10 |
| GAM-40 | 103.24 | 115.91 | 134.74 | 51.78 | **141.80** | 535.70 |
| RF-004 | 0.05 | 0.06 | 0.11 | 0.16 | 0.31 | 0.57 |
| RF-008 | 0.06 | 0.07 | 0.12 | 0.18 | 0.33 | 0.60 |
| RF-016 | 0.07 | 0.08 | 0.14 | 0.20 | 0.36 | 0.65 |
| RF-032 | 0.08 | 0.11 | 0.15 | 0.22 | 0.40 | 0.71 |
| RF-064 | 0.11 | 0.13 | 0.19 | 0.26 | 0.45 | 0.80 |
| RF-128 | 0.15 | 0.18 | 0.25 | 0.32 | 0.53 | 0.89 |
| RF-256 | 0.24 | 0.29 | 0.35 | 0.42 | 0.67 | 1.04 |
| RF-512 | 0.37 | 0.41 | 0.50 | **0.62** | 0.86 | 1.27 |

**Table S-10:**
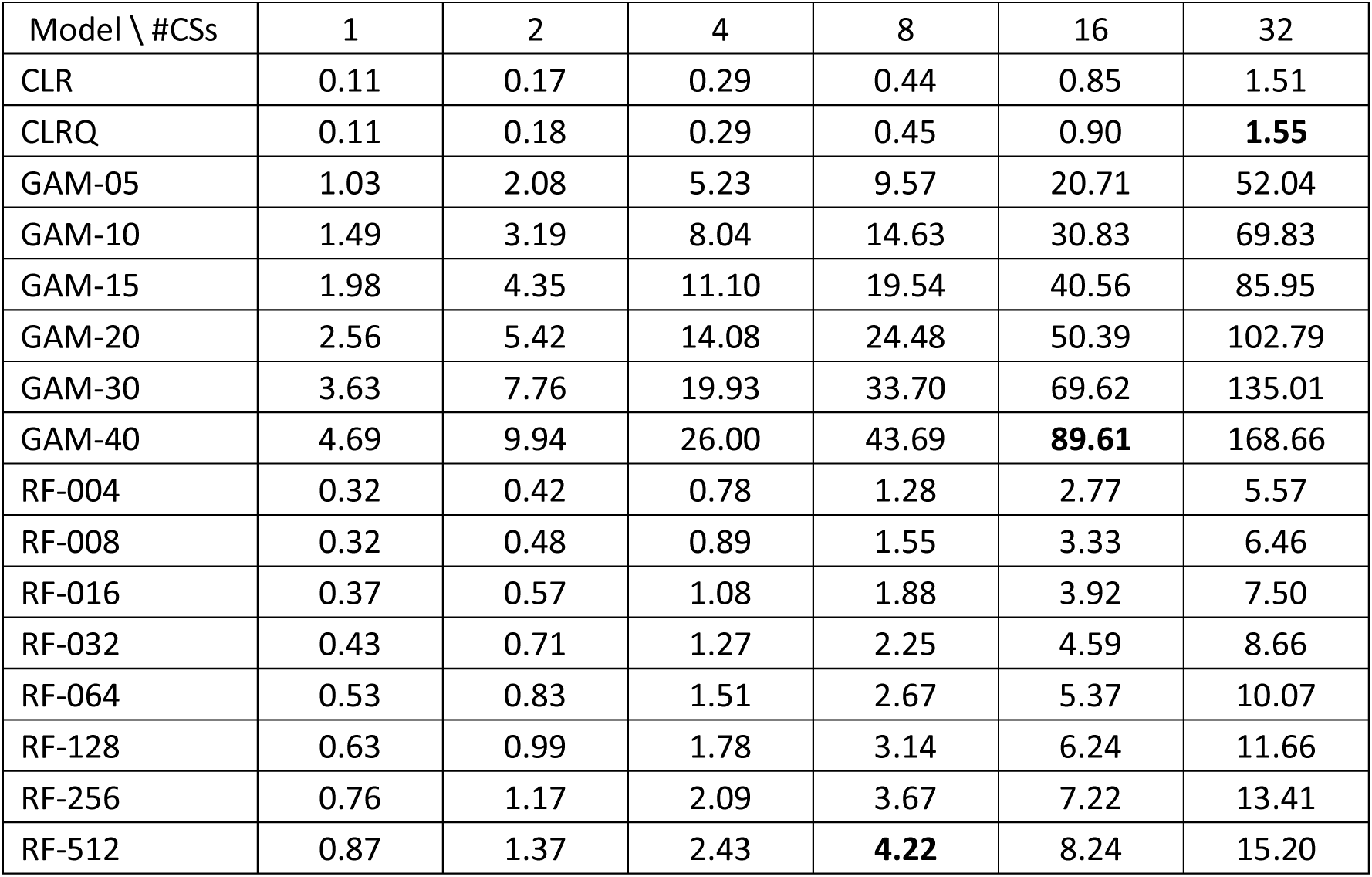
Mean Time to Construct Effect Curve (in seconds)

| Model \ #CSs | 1 | 2 | 4 | 8 | 16 | 32 |
| --- | --- | --- | --- | --- | --- | --- |
| CLR | 0.11 | 0.17 | 0.29 | 0.44 | 0.85 | 1.51 |
| CLRQ | 0.11 | 0.18 | 0.29 | 0.45 | 0.90 | **1.55** |
| GAM-05 | 1.03 | 2.08 | 5.23 | 9.57 | 20.71 | 52.04 |
| GAM-10 | 1.49 | 3.19 | 8.04 | 14.63 | 30.83 | 69.83 |
| GAM-15 | 1.98 | 4.35 | 11.10 | 19.54 | 40.56 | 85.95 |
| GAM-20 | 2.56 | 5.42 | 14.08 | 24.48 | 50.39 | 102.79 |
| GAM-30 | 3.63 | 7.76 | 19.93 | 33.70 | 69.62 | 135.01 |
| GAM-40 | 4.69 | 9.94 | 26.00 | 43.69 | **89.61** | 168.66 |
| RF-004 | 0.32 | 0.42 | 0.78 | 1.28 | 2.77 | 5.57 |
| RF-008 | 0.32 | 0.48 | 0.89 | 1.55 | 3.33 | 6.46 |
| RF-016 | 0.37 | 0.57 | 1.08 | 1.88 | 3.92 | 7.50 |
| RF-032 | 0.43 | 0.71 | 1.27 | 2.25 | 4.59 | 8.66 |
| RF-064 | 0.53 | 0.83 | 1.51 | 2.67 | 5.37 | 10.07 |
| RF-128 | 0.63 | 0.99 | 1.78 | 3.14 | 6.24 | 11.66 |
| RF-256 | 0.76 | 1.17 | 2.09 | 3.67 | 7.22 | 13.41 |
| RF-512 | 0.87 | 1.37 | 2.43 | **4.22** | 8.24 | 15.20 |

In light of the longer times required to construct effect curves for more complex models, we consider an alternative strategy for constructing effect curves, using wolf data (Barry, 2020) as an example. Rather than using all of the nearly 400,000 available combinations, we paired values of a given covariate with 10,000 randomly sampled combinations of the other covariates. The resulting effect curves are shown in Figure S-19. For comparison, we reproduce the version of the effect curves that incorporates all available combinations (reproduced here as Figure S-20, shown in the main paper as Fig. 6). The two sets of plots are very similar to each other, suggesting that sampling available combinations provides a valid and efficient alternative for constructing effect curves.

**Figure S-19:**
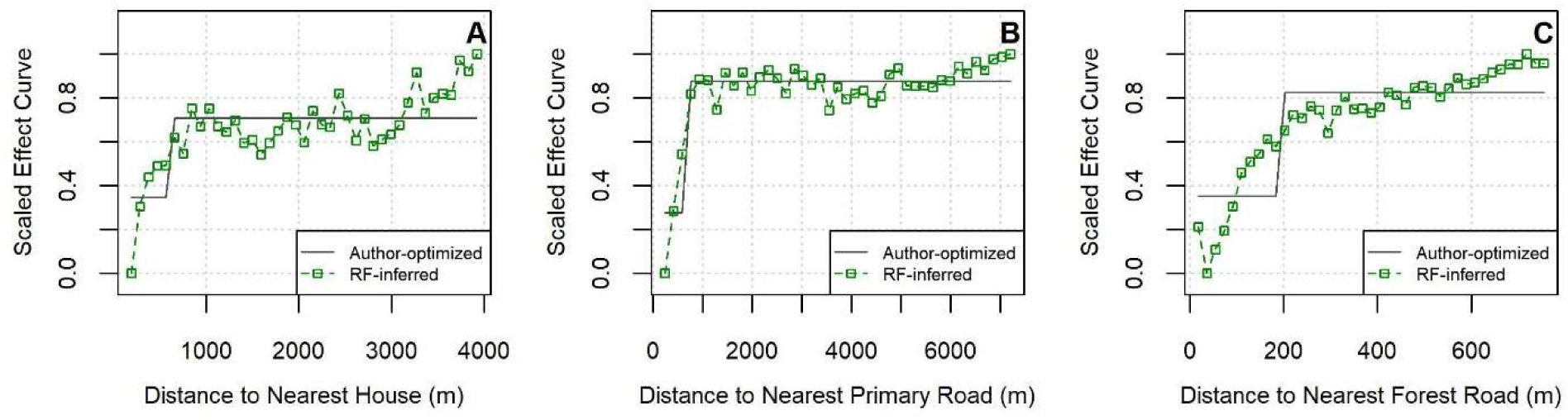
Effect Curves Inferred for Step Selection Covariates for Wolf Tracks (Evaluated with 10,000 Sampled Combinations)

**Figure S-20:**
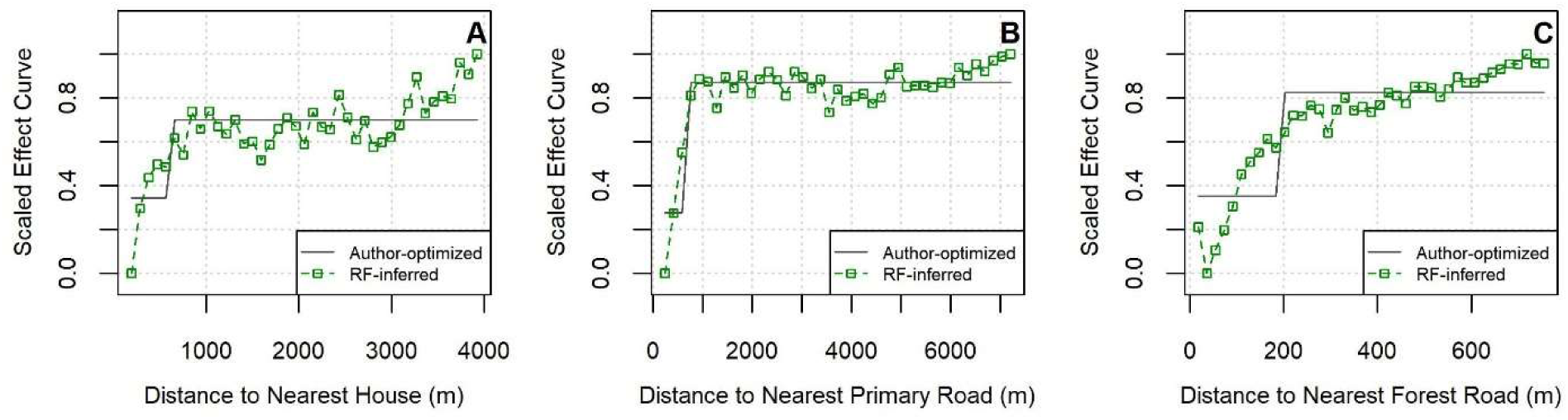
Effect Curves Inferred for Step Selection Covariates for Wolf Tracks (Constructed with 512 Observed Steps and 512 Comparison Steps per Tree, Evaluated with All Available Combinations)

Building on our earlier discussion about the number of CSs and OSs used to train each RF tree (Section 4), we also experimented with including additional data when fitting RFs to these wolf data. Rather than using 512 OSs and 512 OSs per tree (as we used with the simulated tracks in the paper), we set the numbers of OSs and CSs used to train each tree as the number of available OSs. This approach mimics our findings from the paper in a different way. In the paper, we found that the best RF had trees that were trained with an amount of step data that matched the number of available OSs. For the wolf data, this amounts to randomly sampling 43,946 OSs and CSs to train each tree. The effect curves constructed from these expanded models are shown in Figure S-21. These curves reflect the same step-up or ramp-up trends assumed by Barry et al. (2020) or found by our earlier analysis. At the same time, the effect curves from the expanded models appear to contain more noise, perhaps suggesting overfitting.

**Figure S-21:**
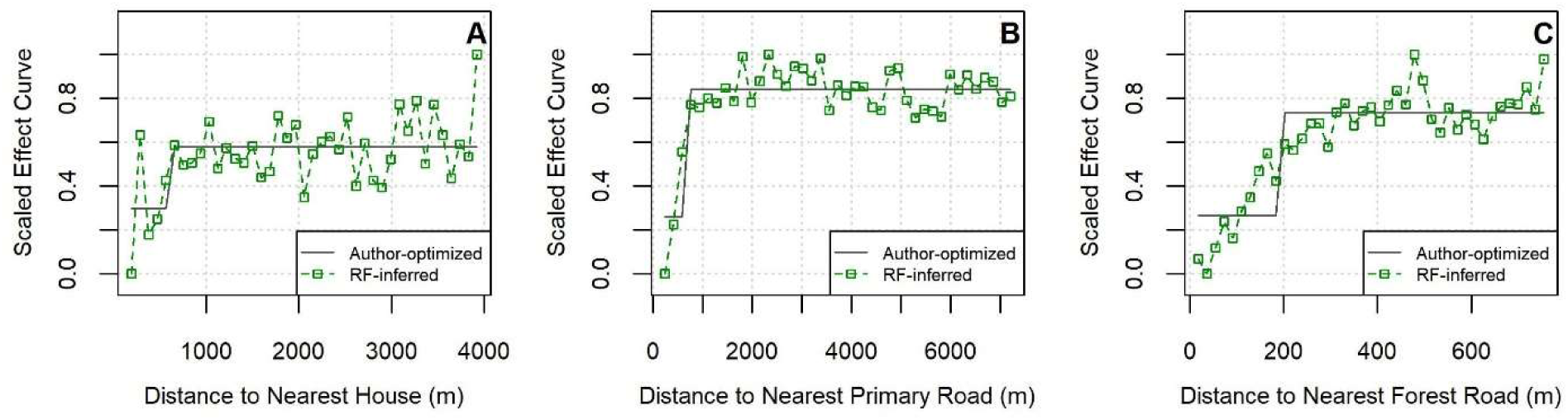
Effect Curves Inferred for Step Selection Covariates for Wolf Tracks (Constructed with Additional Step Data and Evaluated with 10,000 Sampled Combinations)

### S-8. Acronyms

The follow acronyms appear in the main paper or in this supplement.

CLR: Conditional logistic regression
CLRQ: Conditional logistic regression with quadratic term
CS: Comparison step
df: Degrees of freedom
GAM: Generalized additive model
ML: Machine learning
MPS: Milestone proximity score
NDVI: Normalized difference vegetation index
OS: Observed step
PDP: Partial dependence plot
RF: Random forest
RSF: Resource selection function
RTA: Relative turning angle
SL: Step length
SMRF: Semi-matched random forest
SSF: Step selection function
WSR: Wilcoxon signed rank

